# Conformational Priming of RepA-WH1 for Functional Amyloid Conversion Detected by NMR Spectroscopy

**DOI:** 10.1101/612135

**Authors:** David Pantoja-Uceda, Javier Oroz, Cristina Fernández, Eva de Alba, Rafael Giraldo, Douglas V. Laurents

## Abstract

How proteins with a stable globular fold acquire the amyloid state is still largely unknown. RepA is a versatile plasmidic DNA binding protein, functional either as a transcriptional repressor or as an initiator or inhibitor of DNA replication, the latter through the assembly of an amyloidogenic oligomer. Its N-terminal domain (WH1) is responsible for discrimination between these functional abilities by undergoing hitherto unknown structural changes. Furthermore, when expressed alone, RepA-WH1 behaves as a synthetic prion-like protein causing an amyloid proteinopathy in bacteria. RepA-WH1 is a stable dimer whose conformational dynamics had not been explored. Here we have studied it through NMR {^1^H}-^15^N relaxation and H/D exchange kinetics measurements. The N- and the C- terminal α-helices, which lock the WH1 fold in each subunit of the dimer, as well as an internal amyloidogenic loop, show reduced stability and are partially unfolded in solution. S4-indigo, a small molecule ligand known to interfere with the amyloid assembly of RepA-WH1, binds to and tethers the N-terminal α-helix and a β-hairpin that is involved in dimerization, thus providing evidence for a priming role of fraying ends and dimerization switches in the amyloidogenesis of folded proteins.

## Introduction

Numerous human diseases of great social and economic impact are caused by amyloidogenic proteins. Most frequently, these proteins are intrinsically disordered or contain unstructured regions, which facilitates their conversion into amyloid. In contrast, some pathological amyloids arise from well-folded globular proteins, such as transthyretin^1^ or β2-microglobulin^2^. In these cases, amyloid formation is more complex as it involves a prerequisite unfolding event, whose atomic details are only now being elucidated^3^.

RepA protein regulates in the bacterium *Pseudomonas savastanoi* the replication of the plasmid pPS10 which contains directly repeated sequences (iterons) at its replication origin (*ori*). RepA exists in three forms^4–5^: a dimer, which represses transcription of its coding gene; a meta-stable monomer, which triggers DNA replication; and a functional amyloid, whose role is to block new premature rounds of plasmid replication by steric hindrance of the RepA-bound iterons, a process known as handcuffing ^6,7^. This functional amyloid is later removed by chaperones and/or proteases^8^ to allow plasmid replication to resume at a correct pace. The RepA N-terminal Winged Helix domain (WH1) is responsible for these different functional forms, but the conformational changes involved are still unknown.

*In vitro* dissociation of the dimer, formation of a meta-stable monomer and subsequent amyloid assembly can be triggered by sub-stoichiometric amounts of a specific dsDNA sequence (termed *opsp*) from the plasmid replicon, namely: 5’CATTGACTTGT3’/3’GTAACTGAACA5’ ^5^. Amyloid-like fibrils associate to form single or double threads, whose low resolution structure has been solved by TEM ^9^. Since three or more repetitions of the amyloidogenic segment of RepA WH1 can substitute the N-terminus of Sup35 prion and produce stable phenotypic switching of the *[PSI+]* prion in yeast, RepA-WH1 is considered to be a *bona fide* prion-like protein (aka, prioniod)^10^. This is a valuable and simple model system for studying human functional amyloids, such as CPEB3^11^, as well as pathological amyloids exemplified by TDP-43^12^, because both CPEB3 and TDP-43 structural conversions also appear to be regulated by nucleic acids.

The role of dsDNA in promoting the structural conversion of RepA-WH1 into the functional amyloid form has been supported by the observation that this transformation occurs *in vivo* at the bacterial nucleoid, as revealed by immuno-EM utilizing a conformational antibody specific for amyloid oligomers of the protein^13^. Compounds like tetrasulfonated indigo (hereafter, in short, S4-indigo), which block *in vitro* the interaction between dsDNA and RepA-WH1, defuse the “DNA trigger” and this finding represents an important proof of concept that amyloid formation can be prevented by blocking the interactions between an amyloidogenic protein and allosteric cellular factors that catalyze amyloid conversion^14^.

Mutations in WH1, and the substitution of the second winged helix domain (WH2) in RepA by the fluorescent tracer protein mCherry, can convert RepA-WH1 into bactericidal aggregates^15^. These aggregates show two distinct forms *in vivo*, which have been characterized by microfluidics as distinct strains^16^. One form is a relatively benign, elongated aggregate, whereas the second form is a compact, cytotoxic particle which arrests cell grow^16^. Interestingly, the Hsp70 chaperone DnaK can promote the conversion of the compact, toxic form into the elongated form^16^. It is also fascinating that acidic phospholipids, such as POPG and cardiolipin, can promote the aggregation of RepA-WH1-mCherry into pore-like structures in the membrane^17^. Similar results have been reported for the polypeptides Aβ and α-synuclein, which are implicated in Alzheimer’s disease^18^ and Parkinson’s disease^19^, respectively. RepA-WH1 has been exploited to ascertain the effects of amyloidosis on global physiology and metabolism^20^ in a simple bacterial system. A review on RepA-WH1 as a model cytotoxic prionoid has been published recently^21^.

The 3D structure of the native fold of RepA-WH1, composed of five α-helices and five β-strands, has been solved by X-ray crystallography at high resolution^22^ (PDB code 1HKQ), and provides some clues on the elements primed to form amyloid. In most globular proteins with a combination of α-helical and β-sheet secondary structures, the helices typically pack against the central β-sheets^23^. In contrast, the topology of the RepA-WH1 dimer features a highly unusual peripheral helical subdomain (**Sup. Fig. 1**). High B-factors were reported in the helices 1 and 5, which suggests a high intrinsic mobility. Helices 1, 2 and 5 pack to form a latch closing the domain which is stabilized by hydrophobic contacts between L12, L19 (helix 1); L26, V27, I34 (helix 2); W94 (in a turn between β-strands 3 and 4); and I115, L119 and L122 (α-helix 5). The high proportion of flexible Leu relative to stiffer, β-branched Val or Ile residues could afford conformational malleability in the first and last helices. Note that the last “anchor” of helix 5 to this subdomain is L122. Beyond L122, α-helix 5 extends another eight residues out into solution without mooring. Indeed, in a model of monomeric *E. coli* RepA-WH1, based on the X-ray crystal structure of the homologous monomeric protein RepE54^24^, the second half of α-helix 5 (residues 121-130) forms a loop and a short β-strand, enabling tight packing of domains WH1 and WH2, and the second half of α-helix 1 (residues 15-19) becomes a loop^24^. Considering that the loss of this amount of helical structure is consistent with CD spectral changes observed upon activation^25^, this monomeric structure^24^ can be taken as a reliable proxy for the metastable monomeric form, which is “activated” for DNA replication.

Despite the functional relevance of RepA-WH1, not much is known about its solution structure and dynamics. This could be key for understanding how structural elements are primed for amyloid conversion, how DnaK chaperones bind to and eventually break down the amyloid^16, 26^ and how certain molecules modulate the equilibria among distinct conformers. For many years, RepA-WH1 has proved to be intractable to study by NMR, due to its rather large size (a dimer with 134 residues per subunit) and, furthermore, a dimer-to-tetramer equilibrium under quasi-physiological conditions leads to severe line broadening for many key residues. Here, taking advantage of RepA-WH1 extraordinary conformational stability, these difficulties have been overcome by using high temperature, low pH and low ionic strength conditions, as well as application of the TROSY module^27^, which selects the fine component of ^1^H-^15^N HSQC type correlations at high magnetic field to register a suite of 3D spectra. The analysis of these spectra has afforded the almost complete backbone resonance assignment of the protein. To assess to what extent the elements of secondary structure detected by X-ray crystallographic analysis are populated in solution, a series of NMR based analyses were performed; namely: 1) conformational chemical shifts, 2) ^3^J_HNCA_ coupling constants, 3) ^1^HN temperature coefficients, 4) H/D exchange. Finally, ^15^N-^1^H NMR relaxation experiments were recorded to determine the backbone dynamics of the protein^28^.

The assigned RepA-WH1 ^1^H-^15^N HSQC NMR spectrum also represents a valuable tool to map intermolecular interactions. Thus, the titration of RepA-WH1 with S4-indigo, a known inhibitor of the amyloidogenesis of the protein’s *in vitro*, monitored here has revealed that S4-indigo binding to the *opsp* DNA recognition site actually partially unlocks the dimerization interface. In addition, a novel S4-indigo recognition site between α-helix 1 and β-strand 2 in each subunit has been identified. On the basis of this discovery, we advance an updated model describing contrasting effects of ligand binding; namely 1) loosening the dimer interface, 2) sterically blocking the *opsp* DNA binding site to impede DNA binding from springing amyloidogenesis and 3) decreasing the tendency to form amyloid by tethering amyloidogenic segments to the more stable protein structural core.

## Materials & Methods

Buffer solutions were prepared in 85% mQ H_2_O / 15% D_2_O or 100 % D_2_O, (Euroisotop), 99.9 % atom D). Sodium-4,4 dimethyl-4-silapentane-1-sulfonate (DSS, from Stohler Isotope Chemicals) was added to samples to a final concentration of 0.1 mM and the intense upfield singlet arising from its trimethyl moiety was used as the ^1^H chemical shift reference. Deuterated acetic acid (d_4_, 99.5 % atom D) was purchased from Sigma/Aldrich.

### RepA-WH1 labeling and purification

RepA-WH1 was expressed from the pRG-H_6_-WH1(wt) plasmid^4–5^ in the *E. coli* BL21(DE3) strain carrying the helper plasmid pRIL-*lacI*^*q*^. Pre-inocula were grown on four fresh M9+Ap_100_+Cm_30_ agar plates incubated overnight at 37 °C. Cells were then rubbed and inoculated into 2 L of M9 media containing [^15^N] NH_4_Cl and [^13^C] glucose, supplemented with Ap_100_, and grown at 37 °C to OD_600nm_ = 0.5. Then, IPTG was supplied to 1.0 mM and the culture further incubated at 37 °C for 8 h. Final cell yields range from 4.5-5.0 g (wet cell mass). Purification then proceeded as described ^4–5^. Briefly, it consisted in obtaining a clarified cell lysate through sonication and ultracentrifugation, followed by Ni^2+^-IMAC, removal of the His_6_-tag with thrombin (which leaves an extra Gly-Ser dipeptide at the N-terminus), SP-sepharose chromatography and Amicon concentration. Homogeneous RepA-WH1 preparations were obtained, with an average yield of 40 mg of pure protein per liter of culture. MALDI-TOF mass spectrometry confirmed a nearly complete ^15^N/^13^C labeling. The protein’s concentration in solution (as mononer) was estimated by absorbance using an extinction coefficient at 280 nm of 13,000 M^−1^ cm^−1^.

#### Synthesis of RepA-WH1 operator dsDNA (*opsp*)

The dsDNA sequence (5’-CATTCACTTGT-3’/ 3’GTAACTGAACA5’) used in the binding studies was synthesized, purified and annealed as indicated^2^. Buffer exchange was performed by gel filtration using a PD-10 column (GE Healthcare).

#### NMR assignment and analysis

All spectra necessary for NMR assignment and dynamics assessment were performed on a Bruker AV 800 MHz (^1^H) NMR spectrometer equipped with a triple-resonance TCI cryoprobe, and an active shielded Z-gradient coil, in 1.0 mM deuterated acetic acid, 85% Milli-Q H_2_O, 15% D_2_O (Euroisotop), pH 4.0 at 50.0 °C, except for one confirmatory dynamics study performed on a Bruker 600 MHz spectrometer fitted with a cryoprobe and Z-gradients. Proton chemical shifts were referenced to the internal reference DSS, and ^15^N and ^13^C chemical shifts were referenced indirectly using gyromagnetic ratios of ^15^N:^1^H and ^13^C:^1^H as recommended^29^.

The strategy for resonance assignment was based on the acquisition of the following 2D and 3D experiments incorporating TROSY modules: 2D ^1^H-^15^N HSQC, 3D HNCO, HNcaCO, HNCA, HNcoCA, CBCAcoNH, HNCACB and hCCCcoNH (15 ms mixing time). A 3D HNHA experiment was also performed to assign the ^1^Hα and to calculate de ^3^JHNHα coupling constants^30^.

To identify secondary structure, we utilized the RepA-WH1 amino acid sequence and the ^13^CO, ^13^Cα, ^13^Cβ and ^15^N chemical shifts values as input to perform conformational chemical shift analysis using the web server CSI 3.0^31^. The same data were employed to determine the major *cis/trans* conformational state of the X-Pro bonds using the program Promega^32^.

Spectra were processed with Topspin (Bruker Biospin, Karlsruhe, Germany) or NMRPipe^33^ and analyzed with SPARKY^34^ and NMRView^35^. The backbone and side chain ^1^H, ^13^C and ^15^N chemical shifts have been deposited in the BioMagResBank (http://www.bmrb.wisc.edu/) under accession number 27837. To assign 2D ^1^H-^15^N HSQC under more physiologically relevant conditions; namely 5.0 mM MgSO_4_, 15.0 mM KH_2_PO_4_/K_2_HPO_4_, pH 6.1 and 37.0 °C, a series of ^1^H-^15^N HSQC spectra were recorded on ^13^C,^15^N-RepA-WH1 sample initially at pH 4.0 in 1.0 mM deuterated acetic acid at 50.0 °C, to which small aliquots of the buffer indicated above were added. Then, additional spectra were recorded at 45.0, 40.0 and 37.0 °C. The assignments were confirmed by recording a 3D HNCO spectrum on the sample in the final conditions.

### ^1^H^15^N relaxation measurements and analysis

Spectra to determine the ^15^N *R*_1_ and *R*_1rho_ values and {^1^H}-^15^N NOE of RepA-WH1 uniformly labeled with ^13^C and ^15^N were acquired using the approach of Kay and co-workers^36^. The spectra in each experiment were recorded in an interleaved fashion. No linear prediction, which can affect peak intensities, was utilized in the data processing. The spectral width was 13 ppm and 32 ppm for ^1^H and ^15^N, respectively. Seven delays (60, 100, 240, 460, 860, 1260, and 1600 ms) were used for *R*_1_ measurements, and a different set of ten delays (8, 16, 24, 32, 48, 76, 116, 224, 300, and 500 ms) was utilized to measure the *R*_1rho_ values. These values and uncertainties were calculated by fitting a single exponential equation to the data. To allow solvent relaxation and ensure maximal development of {^1^H}-^15^N NOEs before acquisition, we followed the recommendations of Renner *et al.*^28^ and used a long recycling delay of ten seconds. Three sets of experiments at 800 MHz were recorded: two at 50.0 °C, with or without a four fold excess of S4-indigo, and one at 37.0 °C without S4-indigo, all in 1.0 mM DAc (d4), pH 4.0. Two additional sets of experiments: 1) at 600 MHz under these pH 4.0 conditions and 2) at 800 MHz in 5.0 mM MgSO_4_, 15 mM KH_2_PO_4_/K_2_HPO_4_, pH 6.1 at 50.0 °C, showed lower signal/noise ratios and were not included in the analysis.

The *R*_2_ relaxation rates were then calculated using the equation: *R*_2_ = *R*_1_ρ – *R*_1_ (cos θ)^2^ / (sin θ)^2^, where θ = arctan (1700 Hz / δ ^15^N peak – δ ^15^N spectral center). Relaxation rates were calculated via least-squares fitting of an exponential function (I_t_ = I_0_*e^−kt^ + I_∞_) to peak intensities, using the rate analysis routine of the Java version of NMRView as previously described^37^. For non-overlapped signals, the heteronuclear NOEs were calculated from the ratio of crosspeak intensities in spectra collected with and without amide proton saturation during the recycle delay. Uncertainties in peak heights were determined from the standard deviation of the intensity distribution in signal-less regions of the ^1^H-^15^N HSQC spectra. We estimated the overall correlation time, τ_c_, from the ratio of the mean values of *T*_1_ and *T*_2_ as τ_c_ ≃ 1 / (4 *π* * ^15^N frequency in Hz) * ((6* R_2_/ R_1_) – 7))^½^ which was derived from eqn. 8 of Kay *et al.* ^38^, that takes into account J(o) and J(w) spectral densities and discounts terms of higher frequencies from a subset of residues with little internal motion and no significant exchange broadening. This subset excluded residues with {^1^H}-^15^N NOE ratios below 0.65 and also residues with *T*_2_ values lower than the average minus one standard deviation, unless their corresponding *T*_1_ values were larger than the average plus one standard deviation^39^. The resulting τc value was compared to the correlation time of RepA-WH1 predicted on the basis of its size and shape in solution, using the HYDRONMR program^40^ and the RepA-WH1 crystal structure (PDB: 1HKQ)^22^. Although τ_c_ is often used to estimate the molecular weight using the empirical equation: MW (in kDa) ≃ 1.494·τ_c_ (in ns) + 1.1187, based on the experimental data of Rossi *et al.* ^41^, we found that this relation does not hold at 50 °C, where the solvent viscosity is significantly lower.

The principal components of the RepA-WH1 inertia tensor were calculated with PDPInertia (A. G. Palmer, III, Columbia University, New York, NY) using the X-ray structure (PDB: 1HKQ)^22^. The diffusion tensor, which describes rotational diffusion anisotropy, was determined by two approaches, namely: the r2r1_diffusion^42^, and the quadric_diffusion programs^43^ written by A. G. Palmer, III, (Columbia University). The calculations were successful after using the errors in *T*_1_ and *T*_2_ estimated by Monte Carlo simulations. The ^15^N relaxation was analyzed assuming dipolar coupling with the directly attached proton (bond length = 1.02 Å), and a contribution from the ^15^N chemical shift anisotropy evaluated as −160 ppm. Relaxation data were fitted to the Lipari and Szabo model^44^ using the TENSOR^45^ and FAST-Modelfree programs^46^; the latter interfaces with MODELFREE version 4.2^47^. Five models of internal motion were evaluated for each amide ^1^H-^15^N pair: (i) S^2^, (ii) S^2^ and τ_e_, (iii) S^2^ and *R*_ex_, (iv) S^2^, τ_e_, and *R*_ex_, and (v) S^2^, Sf^2^, and τ_e_, where S^2^ is the generalized order parameter, τ_e_ is the effective internal correlation time, *R*_ex_ is the exchange contribution to transverse relaxation, and Sf^2^ is related to the amplitude of the fast internal motions.

### Temperature coefficients

The ^1^HN temperature coefficients, (Δ δ ^1^H / Δ T), were measured by comparing the ^1^H chemical shift values in TROSY ^1^H-^15^N HSQC spectra acquired at 40.0 °C and 50.0 °C. To facilitate the assignment of peaks at 40.0 °C, and to corroborate the linearity of the temperature coefficients, an additional TROSY ^1^H-^15^N HSQC spectrum was recorded at 45.0 °C.

### Hydrogen/deuterium exchange

H/D exchange was started by transfer of a thawed ^13^C,^15^N RepA-WH1 sample into buffer containing 15 mM KH_2_PO_4_/K_2_HPO_4_, 5 mM MgSO_4_ with 100% D_2_O using a PD-10 gel filtration column that had been pre-equilibrated with this buffer. H/D exchange was monitored by recording a series of 38 ^1^H-^15^N TROSY-HSQC spectra (matrix size: 2k ^1^H, 256 ^15^N, number of scans: 12, sweep width: 13 ppm ^1^H, 32 ppm ^15^N, ^13^C-decoupled, transformed without linear prediction) over a period of 2½ months. The pH* (*i.e.* the pH reading without correction for the deuterium isotope effect) of this sample was 5.69. These experiments were carried at a somewhat lower temperature of 37.0 °C. This allowed the exchange of more amide groups to be monitored relative to 50.0 °C. The decrease in ^1^H-^15^N during the course of exchange was quantified using Dynamics Center 2.5.3 (Bruker Biospin). The same program was utilized to fit a single exponential decay function to the data to obtain the experimental exchange rates (k_exp_). The identity of the exchanging peaks was confirmed by recording a 3D HNCO spectrum on a duplicate sample and then comparing the ^1^HN, ^15^N and ^13^CO chemical shifts.

The intrinsic coil exchange rates (k_coil_) for backbone amide groups were calculated with the parameters reported by Bai, Englander and colleagues^48^ using the on-line program Sphere developed by Y. Z. Zhang and H. Roder^49^. The program’s default parameters were utilized, except the pKa values, which were substituted by the values reported by Pace and coworkers^50^. The Trp 94 NHε coil rate was calculated using previously reported parameters for 5.0 °C^48^ and an activation enthalpy of 17 kcal/mol to extrapolate this coil rate at 37.0 °C. The protection factor (PF) for exchange for each measurable HN group was calculated as ratio of the k_coil_/ k_exp_. The minimal PF that can be detected under our experimental conditions is about 1000, which means that an H-bond must be present more than 99.9% of the time for protection to be observed. The uncertainties in the PF values were calculated by propagating the uncertainties in the fitted experimental rates at the 2σ (95%) confidence level. Under certain conditions, where the EXII exchange mechanism dominates, the per-residue conformational stability (ΔG_HX_) may be calculated as ΔG_HX_ = RT ln (PF). However, the validation of this approach requires knowledge of the protein’s folding and unfolding rates, which are currently unknown. Therefore, the analysis of the H/D exchange experiments is limited here to the level of protection factors.

### RepA-WH1 binding to S4-indigo monitored by NMR

The titration of ^13^C,^15^N RepA-WH1 with S4-indigo (potassium indigotetrasulfonate, K_4_C_16_H_6_N_2_S_4_O_14_, TCI Europe) was followed by TROSY ^1^H-15N HSQC spectra and monitoring changes in the ^1^H-^15^N chemical shifts. Chemical shift perturbations (CSP, Fig. 4A) were calculated using the following formula: CSP (ppm) = [(δ^1^H_bound_ - δ^1^H_free_)^2^ - (δ^15^N_bound_ - δ^15^N_free_) / 5)^2^]^½^ Three independent titrations were performed reaching final S4-indigo : ^13^C,^15^N RepA-WH1 dimer ratios of 2:1, 4:1 and 7:1. To assess the role of Arg side chains in RepA-WH1 binding S4-indigo, the lowest of those titration ratios was used and additional TROSY ^1^H-^15^N HSQC spectra were recorded over the region between 75 ppm and 95 ppm (δ ^15^N) to observe Arg ^1^Hε^15^Nε resonances. Some aggregation was observed when higher ratios of S4-indigo were explored. The assignment of the ^1^H-^15^N resonances was confirmed by recording 3D HNCO spectra at the titration end point and comparing the ^13^CO δ values. For the experiments with S4-indigo: ^13^C,^15^N RepA-WH1 ratios of 4:1 and 7:1, the effects of the ligand on protein conformation and dynamics were characterized by recording 3D HNHA spectra to determine ^3^J_HNHA_ coupling constants, measuring the ^1^HN temperature coefficients using three TROSY ^1^H-^15^N HSQC spectra at 40 °C, 45 °C and 50 °C, and registering the hNOE ratio.

**Fig. 4.**
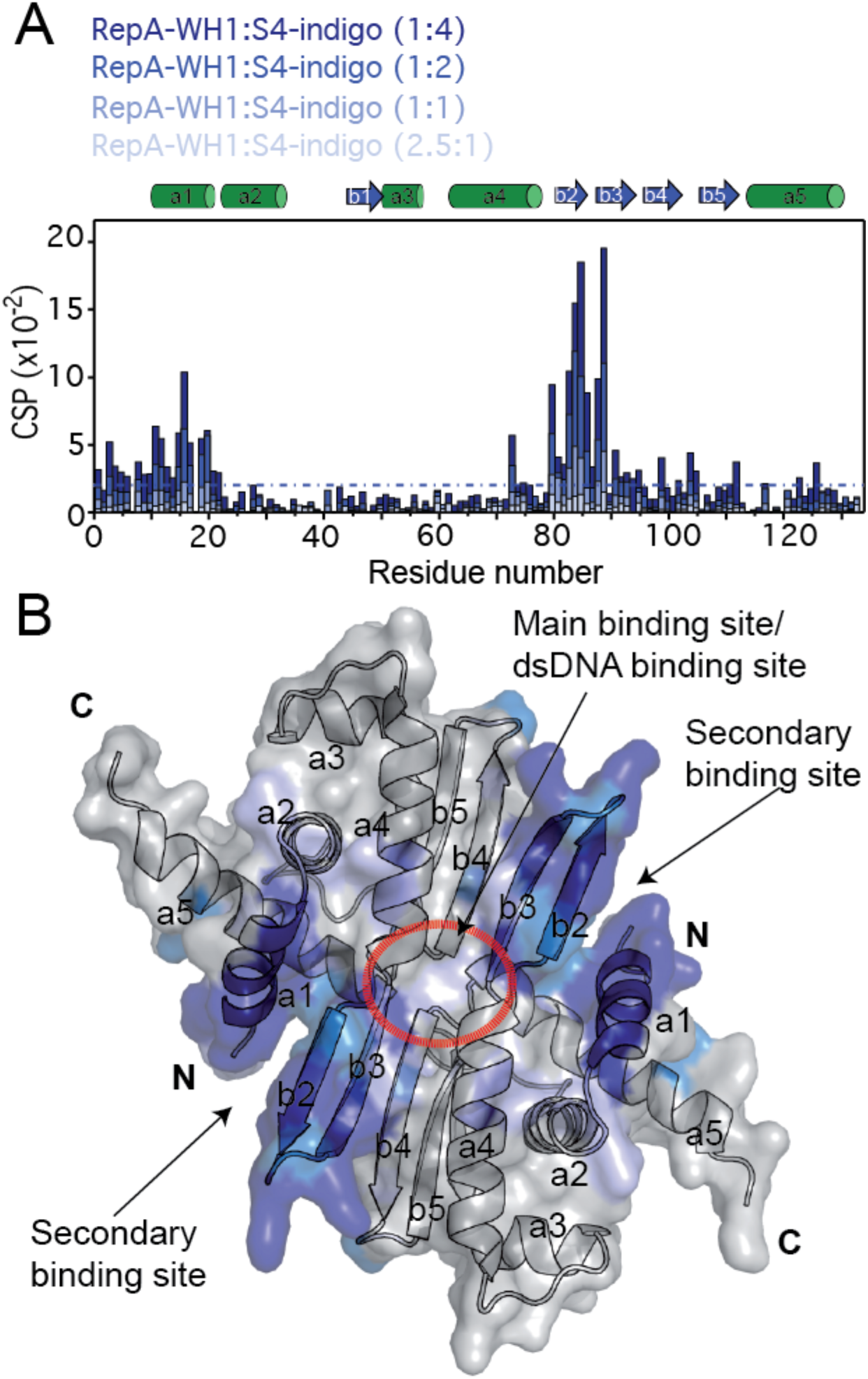
RepA-WH1 / S4-indigo Internation. **A.** Titration of RepA-WH1 with S4-indigo produces ^1^H-^15^N chemical shift changes. Perturbations higher than the dot-dash line are signficant. **B.** Location of binding sites as mapped onto the RepA-WH1 (PDB 1HKQ) structure.

Finally, the amide HN hydrogen bonding of ^13^C,^15^N RepA-WH1 in the presence of a four fold excess of S4-indigo was probed by H/D exchange at 37 °C. This was carried out by dissolving a lyophilized sample of ^13^C,^15^N-RepA-WH1 with four equivalents of S4-indigo in D_2_O and monitoring a series of 2D ^1^H,^15^N HSQC spectra at 37.0 °C in 1.0 mM DAc, pH* 3.6. For comparison, an additional H/D exchange experiment in the absence of S4-indigo was performed at pH* 3.6 in 1.0 mM DAc (d4), 37.0 °C. We also attempted to measure H/D exchange in 15.0 mM KH_2_PO_4_/K_2_HPO_4_, 5.0 mM MgSO_4_, pH* 5.7 to match the conditions of the original H/D exchange experiment described in the preceding section. For the pH* 5.7 experiments, transfer to the deuterated buffer with S4-indigo was attempted both by dissolving lyophillized protein in buffer of by gel filtration with a short PD-10 column. However, both these experiments with S4-indigo at pH* 5.7 failed to yield observable protein signals.

The docking program HADDOCK (high ambiguity-driven protein-protein docking) version 2.2 was used to model the complexes between RepA-WH1 and S4-indigo ^51^. The restraints used for the interaction were generated from the chemical shift perturbations analysis. For RepA-WH1, the crystral structure (PDB: 1HKQ) was used. Active ambiguous interaction restraints (AIRs) included residues whose CSP values are > 0.06 ppm; namely: N10, K11, E14, S15, S16, T18, D79, R82, Y83, V84, K85, K87 and V88. Passive residues were L12, I80, R81, G86, E90, V98, G103, F111 and E125; their CSP values are between 0.03 and 0.06 ppm. The parameter and topology files of indigo-S4 were generate with the JME editor using the PRODGR server ^52^. A total of 500 structures were sampled during the rigid body docking. During the final iteration, the docking models were refined in explicit water. The 50 lowest energy structures were clustered on the basis of the pairwise ligand interface RMSD matrix, using a 2.5 Å cut-off with a minimum of four members per cluster. Finally, an additional round of HADDOCK calculations were conducted using the CSPs plus the H-bond restrictions for the indigo-S4 sulfontate to Q8 side chain H_2_Nε group.

## Results

### Very low ionic strength is key to obtain high quality RepA-WH1 NMR spectra

Multiple solution conditions were surveyed to identify those yielding NMR spectra suitable for analysis. Quasi-physiological conditions (pH 6.1 in 5.0 mM MgSO_4_ and 15.0 mM KH_2_PO_4_/K_2_HPO_4_, 37 °C) produced good quality 2D ^1^H-^15^N spectra. Nevertheless, in these conditions many of the expected peaks were missing in less sensitive 3D spectra (*i.e.* all except HNCO and HNCA). Using analytical ultracentrifugation, RepA-WH1 was found to exist as a dimer-to-tetramer equilibrium under these conditions. The line broadening of NMR peaks due to conformational exchange upon tetramer formation and dissociation, coupled to the already large size of the RepA-WH1 dimer (134 residues x 2) would well account for the patchy quality of the 3D spectra under these conditions. Cooling to 30 °C or heating to 60 °C did not improve the sample’s spectral quality and increasing the ionic strength worsened it. Applying the TROSY module led to some but insufficient improvement. Considering that harsher solution conditions could destabilize the tetramer relative to the dimer, and the discovery by Song’s laboratory that very low ionic strength conditions at lower pH can often improve NMR spectral quality^53^, which we have recently corroborated^54^, additional spectra were recorded at pH 4.0 in 1 mM deuterated acetic acid at 50.0 °C. These conditions, when combined with the TROSY module, did yield very good quality 2D and 3D spectra. Whereas the signal strength still notably varies along the sequence in these spectra, the resonances were intense enough to follow interresidual connectivities. After assigning RepA-WH1 under these conditions, a series of 2D ^1^H-^15^N HSQC spectra were used to transfer these assignments to quasi-physiological conditions. The assigned HSQC spectrum obtained at 50.0 °C in 1.0 mM deuterated acetic acid, pH 4.0, is shown in Fig. 1A. The ^1^HN, ^1^Hα, ^13^Cα, ^13^CO backbone and ^13^Cβ resonances are completely assigned except for the ^15^N of Pro residues, and the ^1^HN and ^15^N resonances of H17 and L119. The assignment data are listed in **Sup. Table 1** and have been submitted to the BMBR database under accession number 27837.

**Table 1.** Average Values and Standard Deviations of the Measured 15N Relaxation Parameters and Overall Fitted Isotropic Correlation Times for RepA-WH1 at 50.0 °C (323.15 K) and 18.8 T

| NMR Parameter | Mean value ± standard deviation<br>All residues (1-132) | Mean value ± standard deviation<br>All residues except N-term. (1-9)<br>& C-term. (129-132) |
| --- | --- | --- |
| $R_1(\text{s}^{-1})$ | $1.0 \pm 0.2$ | $0.98 \pm 0.18$ |
| $R_2(\text{s}^{-1})$ | $17.8 \pm 4.6$ | $19.2 \pm 2.1$ |
| $\{^1\text{H}\}\text{-}^{15}\text{N}$ NOE | $0.65 \pm 0.28$ | $0.72 \pm 0.10$ |

**Fig. 1.**
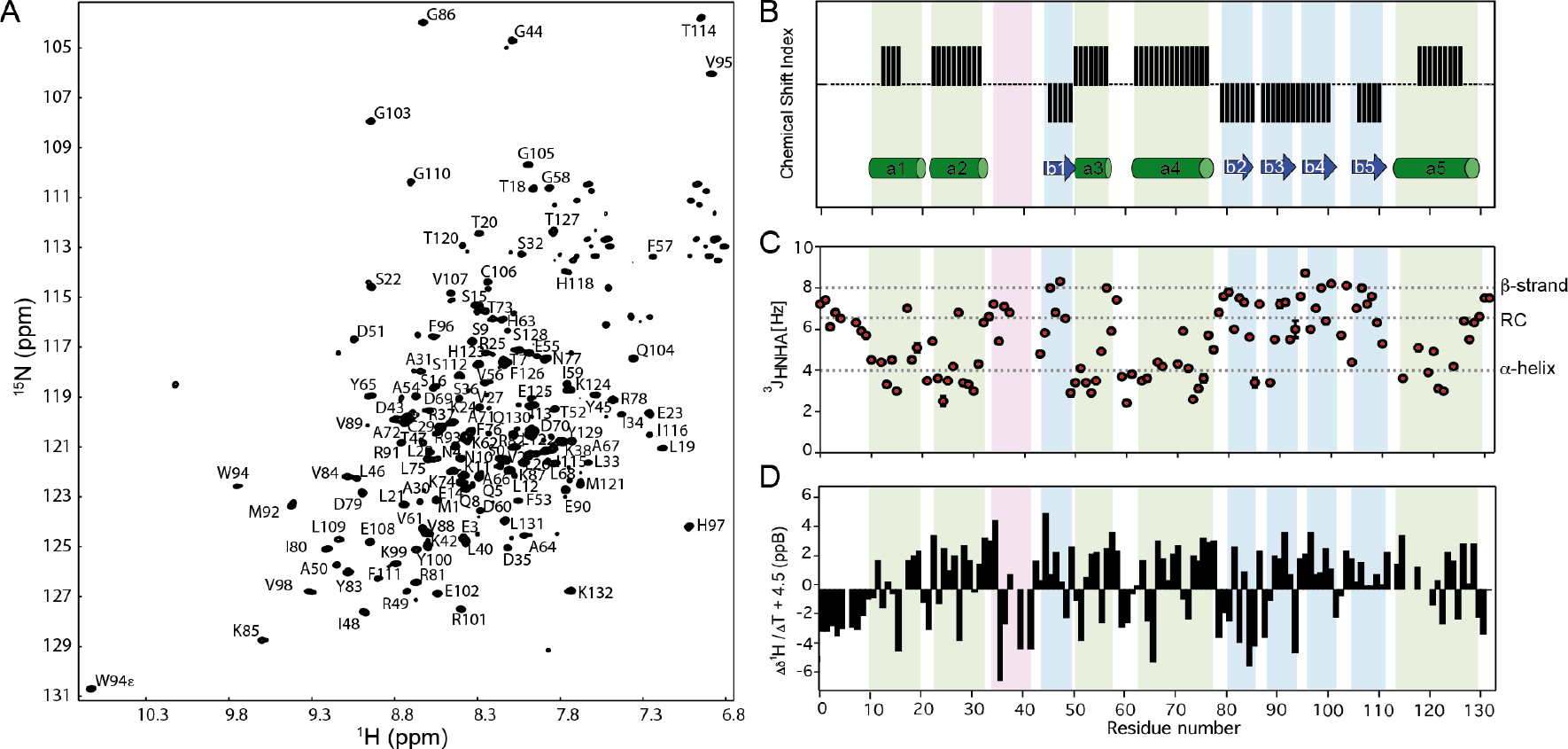
RepA-WH1 Assigned ^1^H-^15^N Spectrum and Solution Secondary Structure. **A.** Assigned ^1^H-^15^N HSQC spectrum of RepA-WH1 recorded in 1.0 mM deuterated acetic acid, pH 4.0, 50.0 °C. **B.** Comparison of the solution secondary structure based on conformational chemical shifts as positive bars (α-helix) and negative bars (β-strands) versus the X-ray crystal structure where α-helices and β-strands are represented by green rectangles and blue arrows, respectively. **C.** J-coupling assessment of secondary structure. ^3^J_HNHA_ values near 4 Hz are characteristic of α-helix, whereas higher values are consistent with coil or β-strand conformations. The contrast between the consistently low ^3^J_HNHA_ values for α-helices 2, 3 and 4 and the higher ^3^J_HNHA_ values near the N- and C-termini of α-helices 1 and 5 is evident. **D.** ^1^HN temperature coefficients to reveal hydrogen bonded HN groups. For clarity, a value of 4.5 ppB / K was added to each data point so that ^1^HNs with temperature coefficients indicative of H-bonding will appear above zero and non H-bonding HN groups will appear < 0. α-helices and β-strands are shaded green and blue, respectively. The amyloidogenic loop is shaded magenta. Whereas the HN groups in the first turn of α-helices generally lack H-bond acceptors and do not form H-bonds, here it can be noted that the proportion of non-hydrogen bonded HN groups is higher in α-helices 1 and 5 than α-helices 2, 3 and 4. The experimental uncertainty in these values is about ± 0.3 ppB·K^−1^.

### NMR spectral analysis reveals incomplete formation of helices 1 and 5 in solution

The structure-induced differences in assigned chemical shifts relative to statistical coil values provide a powerful tool to determine the type and population of elements of secondary structure in solution^55^. Analysis of these “conformational chemical shifts” reveals that the β-strands and α-helices 3 and 4, which form the structural core of RepA-WH1, have a similar extension and position in solution as compared to the X-ray crystal structure. In contrast, in the helical subdomain, α-helices 1 and 5 appear five and eight residues shorter, respectively, in solution than in the crystal structure (Fig. 1B).

RepA-WH1 contains four proline residues, all of whose Xaa-Pro peptide bonds are in the *trans* conformation in the X-ray crystal structure (PDB:1HKQ). The average Pro ^13^Cβ chemical shift values are 27.3 ppm for *cis* Xaa-Pro peptide bonds and 32.1 ppm for *trans* Xaa-Pro peptide bonds. Considering that the ^13^Cβ chemical shift values for Pro 39, 41, 113 and 117 are 32.5, 32.5, 31.9 and 31.2 ppm, respectively, it can be concluded that all four Xaa-Pro peptide bonds are also in *trans* in the folded solution structure. These conclusions have been corroborated by the program PROMEGA. Two Cys residues, C29 and C106, are present in RepA-WH1. In the X-ray crystal structure, the γSH moieties of these Cys ligate two different mercury ions, which were present for phase determination. In solution, the mean ^13^Cβ chemical shifts for reduced and oxidized Cys are 28 and 41 ppm, respectively. Since, the experimental ^13^Cβ chemical shifts for C29 and C106 are 27.1 ppm and 32.4 ppm, respectively, these data indicate that both Cys are reduced under our solution NMR conditions, despite the absence of an added reducing agent. C29 is near the C-terminus of α-helix 2 and C106 lies in β-strand 5. As noted above, these elements of secondary structure appear similar in solution as in the crystal structure, which suggests that the mercury ion binding does not significantly perturb the secondary structure.

J-coupling analysis provides an independent way to assess protein secondary structure in solution ^56^. The mean ^3^J_HNHA_ values for RepA-WH1 at pH 4.0 and 50.0 °C confirm the incomplete formation of helices 1 and 5 and suggest that helices 2, 3 and 4 are completely structured (Fig. 1C).

### Corroboration of the fraying of helices 1 and 5 by ^1^HN temperature coefficients

The ^1^HN temperature coefficients, Δ δ ^1^H / Δ T, for RepA-WH1 were calculated from the ^1^H chemical shift values in TROSY ^1^H-^15^N HSQC spectra recorded at 40 °C and 50 °C and are shown in Fig. 1D. The ^1^H of the first twelve residues or in loops typically show Δ δ ^1^H / Δ T values < −4.5 ppB / K; this is indicative of a lack of H-bonds. In contrast, most ^1^HN in α-helices and β-sheets^57^ and polyproline II helical bundles^58^ show Δ δ ^1^H / Δ T > −4.5 ppB / K, which is consistent with H-bond formation. However, the proportion of residues with Δ δ ^1^H / Δ T > −4.5 ppB / K is less in helices 1 and 5 (Fig. 1D), which suggests that these helices are fraying and lack many H-bonds under these experimental conditions.

### Backbone dynamics from ^15^N relaxation reveal heightened flexibility in RepA-WH1’s amyloidogenic loop

We measured the ^15^N relaxation parameters for RepA-WH1 (**Sup. Fig. 2**) at pH 4.0, 50.0 °C in 1.0 mM DAc (d4). The heteronuclear {^1^H}-^15^N NOE and the longitudinal (T_1_) and transverse (T_2_) relaxation times were measured for 121 of the 134 residues of RepA-WH1 (all except G-1, S0, H17, R25, L40, A66, R93, L119 and F126 whose signals were missing or overlapped, and the four prolines). The average values of the ^15^N relaxation parameters are summarized in Table 1. Several residues deviate from the average, mostly at the disordered N-(M1-S9) and C-termini (S128-K132) and the amyloidogenic loop between α-helix 2 and β-strand 1 (D35-D43), for which small {^1^H}-^15^N NOE ratios (**Sup. Fig. 2**) indicate flexibility on fast time scales (ps to ns). Whereas the signal/noise ratios of their data were lower, additional sets of experiments, recorded at 600 MHz under these pH 4.0 conditions, or at 800 MHz at 37.0 °C, pH 4.0 or at 50.0 °C in 5.0 mM MgSO_4_, 15 mM KH_2_PO_4_/K_2_HPO_4_, pH 6.1 all produced similar results. Due to its superior quality, the data set registered at 50.0 °C, pH 4.0 was analyzed in more detail.

Initially, the value of overall correlation time of 10.76 ns was estimated from the ratio of the mean values of T_1_ and T_2_ (see **Material and Methods**). The principal components of the inertia tensor, calculated for the crystal structure (PDB: 1HKQ), modified to included the disordered N-terminal disordered segment as described below, have relative values of 1.00, 0.75, and 0.50. These values indicate that the shape of the protein deviates from that of a sphere and is best represented as a prolate ellipsoid. In agreement with this finding, the diffusion tensor that best fits the relaxation data was also anisotropic, with different values for the diffusion constants parallel and orthogonal to the long axis of the molecule. Their ratio (D‖/D⊥) equals 1.29 ± 0.03. The final global rotational diffusion correlation time for RepA-WH1 produced by the FAST-Modelfree analysis described in the next paragraph is 11.34 ± 0.05 ns. This value is in good agreement with the correlation time obtained from hydrodynamic calculations for the RepA-WH1 dimer (13.26 ns), which takes into account the decreased solvent viscosity at 50.0 °C.

The dynamic analysis was first performed with the program TENSOR. This analysis revealed that the N- and C-termini and the amyloidogenic loop following α-helix 2 are flexible. However, the TENSOR analysis assigned a large number of residues to complex dynamic models and produced very low experimental uncertainties, which in our experience is rather unusual. Therefore, we proceeded to analyze the data with the FAST-Modelfree program. This program requires a 3D structure, which is lacking for the first seven N-terminal residues. Since the lack of electron density in crystal structures suggests mobility and since the NMR data point to the N-termini being disordered, we modeled structurally this segment as extended coil and grafted it onto the PDB 1HKQ structure using Pymol. Using this approach, relaxation data were analyzed employing the model-free formalism to calculate the corresponding dynamics parameters for the amide ^1^H-^15^N pair of each residue. For most of the residues (61%) analyzed, the ^15^N relaxation parameters could be satisfactorily fit to one of the two simpler models, which describe the internal dynamics of the molecule in terms of a generalized order parameter S^2^ and an effective correlation time τe for fast motions (Fig. 2A,C). For 33% of the residues, it was necessary to include a contribution of slow motions to the transverse relaxation time, on the microsecond to millisecond time scale, which are characterized in terms of conformational exchange rate R_ex_ (Fig. 2B). In other cases (six residues: V2, E3, V6, K38, L131 and K132), the inclusion of the amplitude of internal motions (Sf^2^) was also required to obtain a good fit. Finally, two residues (L21 and G103) were not well fit by any model.

**Fig. 2.**
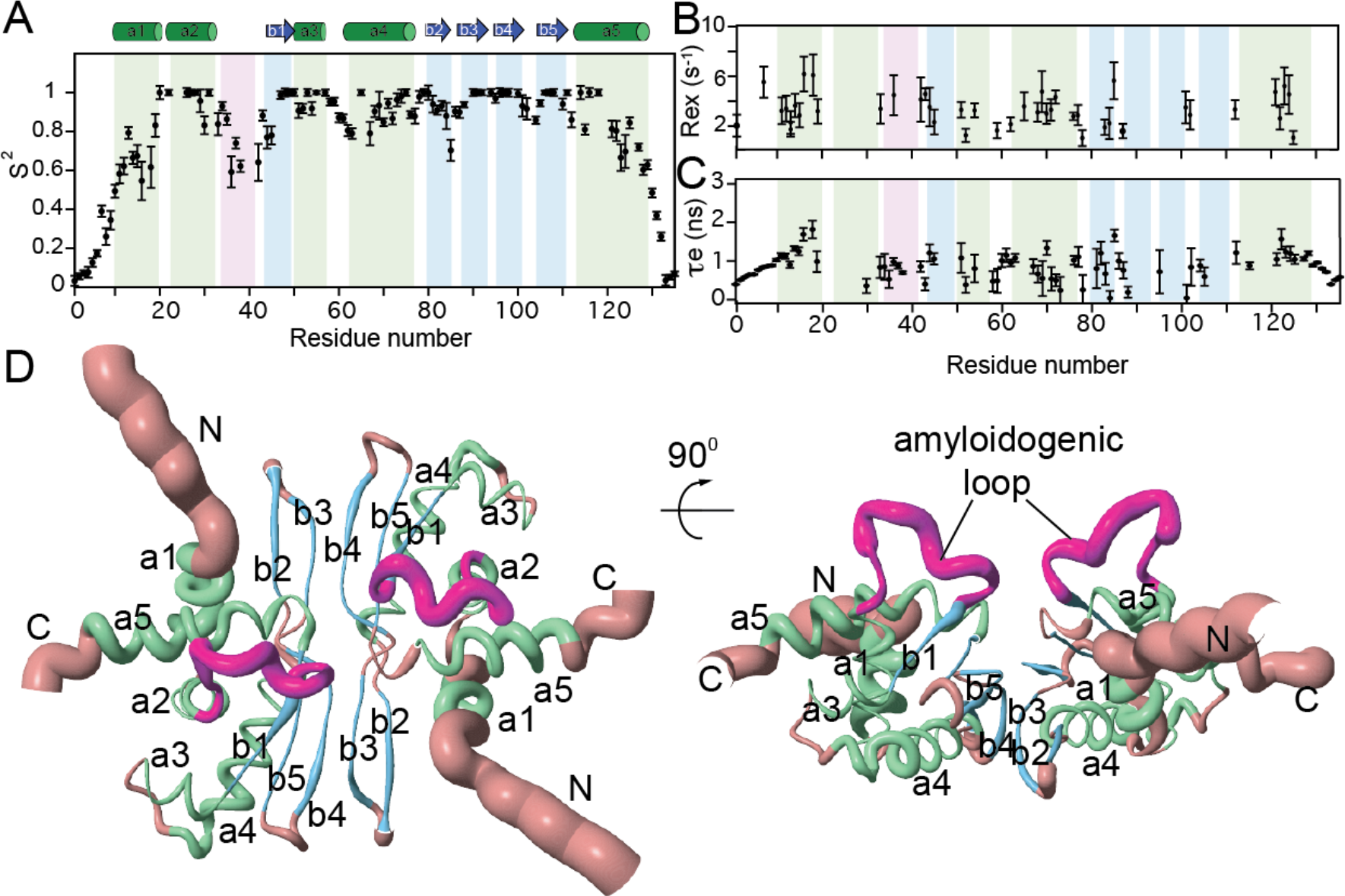
The Amyloidogenic Loop is Flexible on Fast ps/ns and Slow Sec Timescales. **A.** The residue level order parameters (S^2^) **B.** Conformational exchange rates (*R*_ex_) **C.** Effective internal correlation times (τe). In **A**, **B**, and **C**, α-helices are shown in green, β-strands in blue, and the amyloidogenic loop in magenta. **D.** Two views of the RepA-WH1 dimer are shown, with the elements of secondary structure colored as in panel **A-C**, while loops and termini are shaded orange. The width of the tube is proportional to S^2^, so that more dynamic segments, such as the termini and amyloidogenic loop, appear thicker.

The order parameters indicate that the N-terminus and most of α-helix 1 (residues 1-18) and the second half of α-helix 5 and the remaining C-terminal residues (120 − 132) are flexible on fast ps/ns timescales (Fig. 2A, D), in agreement with the lack of secondary structure observed for these regions (Fig. 1). β-strand 2, which is a rather exposed edge strand, and the turn linking β-strands 4 and 5 are slightly flexible.. Remarkably, the amyloidogenic loop linking α-helix 2 and β-strand 1 has order parameters of only 0.60, which indicates that it is rather flexible and susceptible for structural conversion (Fig. 2D). β-strand 2, which is a rather exposed edge strand, and the turn linking β-strands 4 and 5 are also slightly flexible (Fig. 2A, D). In agreement with the dynamics on fast timescales, most of the residues experiencing additional motions on slower timescales are at the termini and in the amyloidogenic loop (Fig. 2B). The remaining elements of secondary structure are quite rigid with order parameters approaching 1.0.

### H/D exchange monitored by NMR reveals low stability of α-helices 1 and 5

The protection against deuterium exchange is afforded by hydrogen bonding and burial on the level of individual residues (Fig. 3). The C-terminal half of helix 4 and β-strands 3, 4 and 5, which form the dimer interface, show the highest protection against exchange. If exchange were dominated by the EXII mechanism, wherein the refolding rate is much faster than the intrinsic H/D exchange rate^59^, the conformational stability of these most stable residues would be approximately 9 kcal/mol. It is notable that the HN groups of residues R91 and V98, which donate intersubunit hydrogen bonds to acceptors in the other monomer, have remarkably high protection factors (Fig. 3C). The side chain indole HNε in W94 shows modest protection against exchange (PF = 1000 ± 300). This group lacks an obvious H-bond acceptor group in the X-ray crystal structure, but it is buried together with the nonpolar sidechains of L12, L19, L26, V27, I34, I115, L119 and L122 to form a small hydrophobic core that stabilizes and orients α-helices 1, 2 and 5. No side chain -CONH_2_ moieties show measurable protection against exchange. α-helix 2 is also protected, but less so on the side that contacts α-helix 5. Protection is rather low in α-helix 3 and β-strand 1 and particularly low in α-helices 1 and 5. This is important because α-helix 5 contacts both α-helix 2 and the successive loop, which is amyloidogenic.

**Fig. 3.**
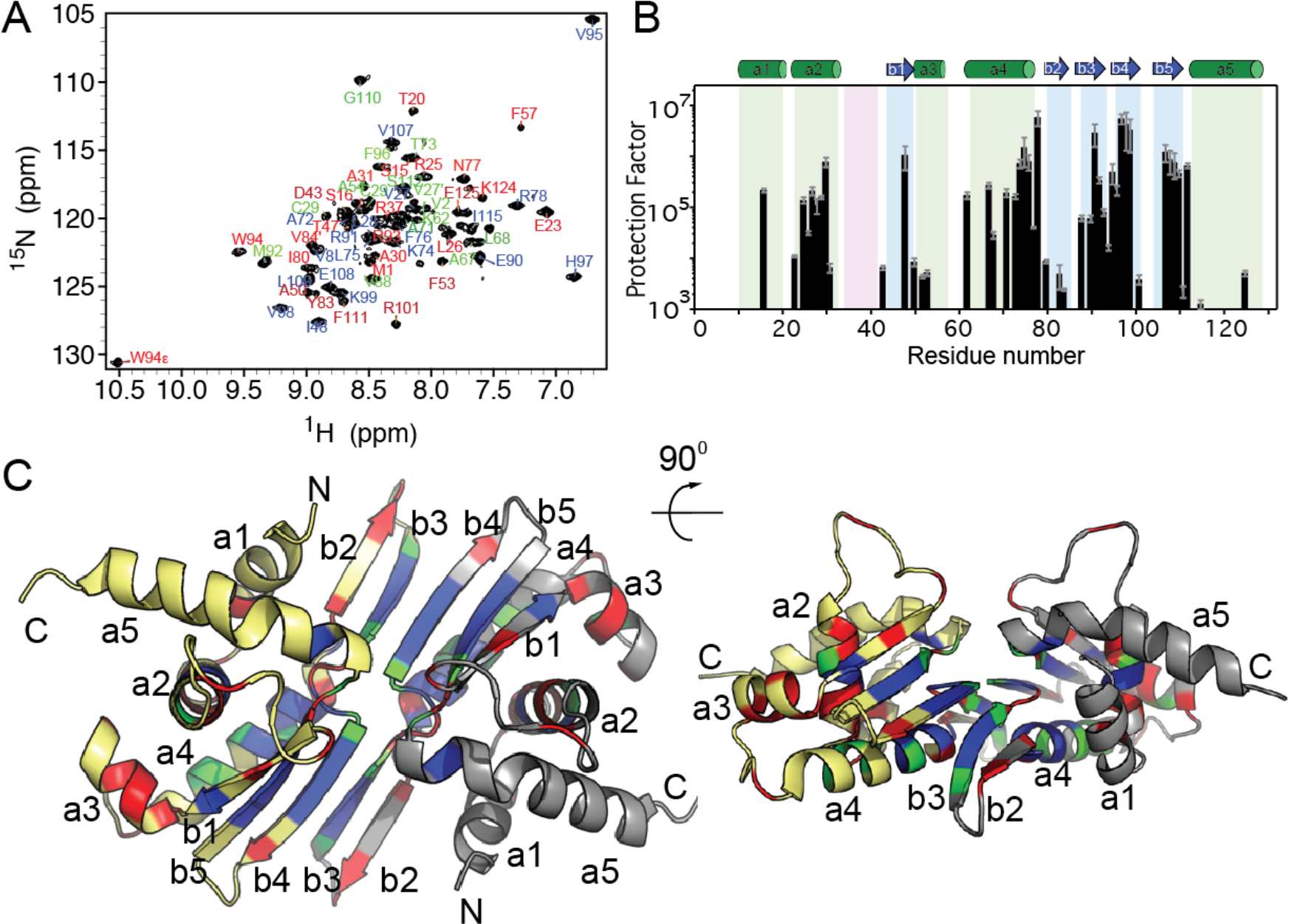
H/D Exchange of RepA-WH1: α-Helices 1 and 5 Show Minimal Protection. **A.** 2D ^1^H-^15^N HSQC spectrum of RepA-WH1 undergoing hydrogen/deuterium exchange. Residues whose HN groups are strongly, moderately and weakly protected against exchange are labeled blue, green or red, respectively. These groups are defined by 2D ^1^H-^15^N HSQC spectra recorded after 72 hours (strong), 15 hours (moderate) and 1 hour (weak) of H/D exchange. Peaks whose identity could not be confirmed by a 3D HNCO spectrum were not labeled. **B.** Per-residue protection factors calculated from H/D Exchange of RepA-WH1 at pH* 5.7, 37.0 °C in 15 mM KH_2_PO_4_, 5 mM MgSO_4_, 100% D_2_O. α-helices, β-strands and the amyloidogenic loop are shaded green, blue and magenta, respectively. Error bars (gray) were calculated by propagation of the uncertainties of the fitted H/D exchange rates **C.** The position of weakly, moderately, and strongly protected backbone HN groups mapped onto the RepA-WH1 structure (PDB 1HKQ; Giraldo *et al.* Nat. Struct. Mol. Biol.). Silver and gold (2^nd^ subunit) colored residues are those whose backbone HN groups lack measurable protection under these conditions.

### RepA-WH1 binding to ligands

The dsDNA 5’-CATTCACTTGT-3’/ 3’-GTAACTGAACA-5’ (*opsp*) binds RepA-WH1, triggering functional amyloid formation. To characterize the dsDNA binding site in RepA-WH1, we followed the dsDNA titration of ^13^C,^15^N RepA-WH1 with 2D ^1^H-^15^N TROSY-HSQC NMR spectra. However, the sample immediately showed signs of precipitation, even at *opsp* dsDNA : protein dimer ratios as low as (1 : 20). This produced a general decrease in the intensity of the protein’s signals, and no more specific information on the interaction could be obtained.

It has been reported that one equivalent of S4-indigo binds strongly (K_D_ = 0.23 μM) to the RepA-WH1 dimer at an region of the subunit interface rich in Arg residues which normally binds to *opsp* dsDNA^14^. In this manner, S4-indigo would compete with dsDNA and prevent RepA-WH1 conformational conversion into amyloid. It was also reported that two or three additional S4-indigo molecules bind a hundred fold more weakly (K_D_ = 23 μM) to the RepA-WH1 dimer^14^, but the locations of these lower affinity binding sites were not determined. In the course of S4-indigo / RepA-WH1 titrations performed here, significant chemical shift or peak intensity changes were observed in the Arg side chain ^1^Hε-^15^Nε resonances (**Sup. Fig. 3**). This is in line with S4-indigo binding to the Arg rich region of the subunit interface, as originally proposed^14^.

Higher ratios of S4 indigo : RepA-WH1 were observed to induce significant changes in the ^1^H-^15^N signals of residues 1-20 (which span the disordered N-terminal region and α-helix 1), and even larger changes in the ^1^H-^15^N resonances of residues 80 – 90, which correspond to β-strands 2 and 3 (Fig. 4). By tethering a weak element of secondary structure (α-helix 1) to more stable elements of secondary structure (β-strands 2 and 3), S4-indigo may induce rigidity and enhance the amount of structure in α-helix 1 as well as α-helix 5, which contacts α-helix 1. To test this hypothesis, both the ^1^HN temperature coefficients (**Sup. Table 2**) and ^3^J_HNHA_ coupling constants (**Sup. Fig. 4A**) were analyzed for evidence of increased hydrogen bonding and augmented formation of helical structure in α-helix 1 and in α-helix 5. Interestingly, several residues in α-helix 1 and the loop preceding α-helix 2 show higher ^1^HN temperature coefficintes (consistent with increased H-bonding) (**Sup. Table 2**). The ^13^CO chemical shift is a better metric of secondary structure than ^1^HN or ^15^N chemical shifts^60^. Although partial precipitation and increased ionic strength eroded NMR spectral quality, it was still possible to acquire good 3D HNCO spectra in the presence of higher concentrations (4x) S4-indigo, and use the assigned ^13^CO chemical shifts to assess whether α-helix 1 became more ordered. However, no ^13^CO chemical shift changes indicative of increased helicity are evident (**Sup. Fig. 4B**).

Changes in {^1^H}-^15^N NOE measurements revealed how S4-indigo binding affects ^13^C,^15^N-RepA-WH1 ps – ns dynamics. The results show that the fast dynamics remain largely similar for α-helix 1 at a four-fold molar excess of S4-indigo relative to RepA-WH1 dimer (Fig. 5A).

**Fig. 5.**
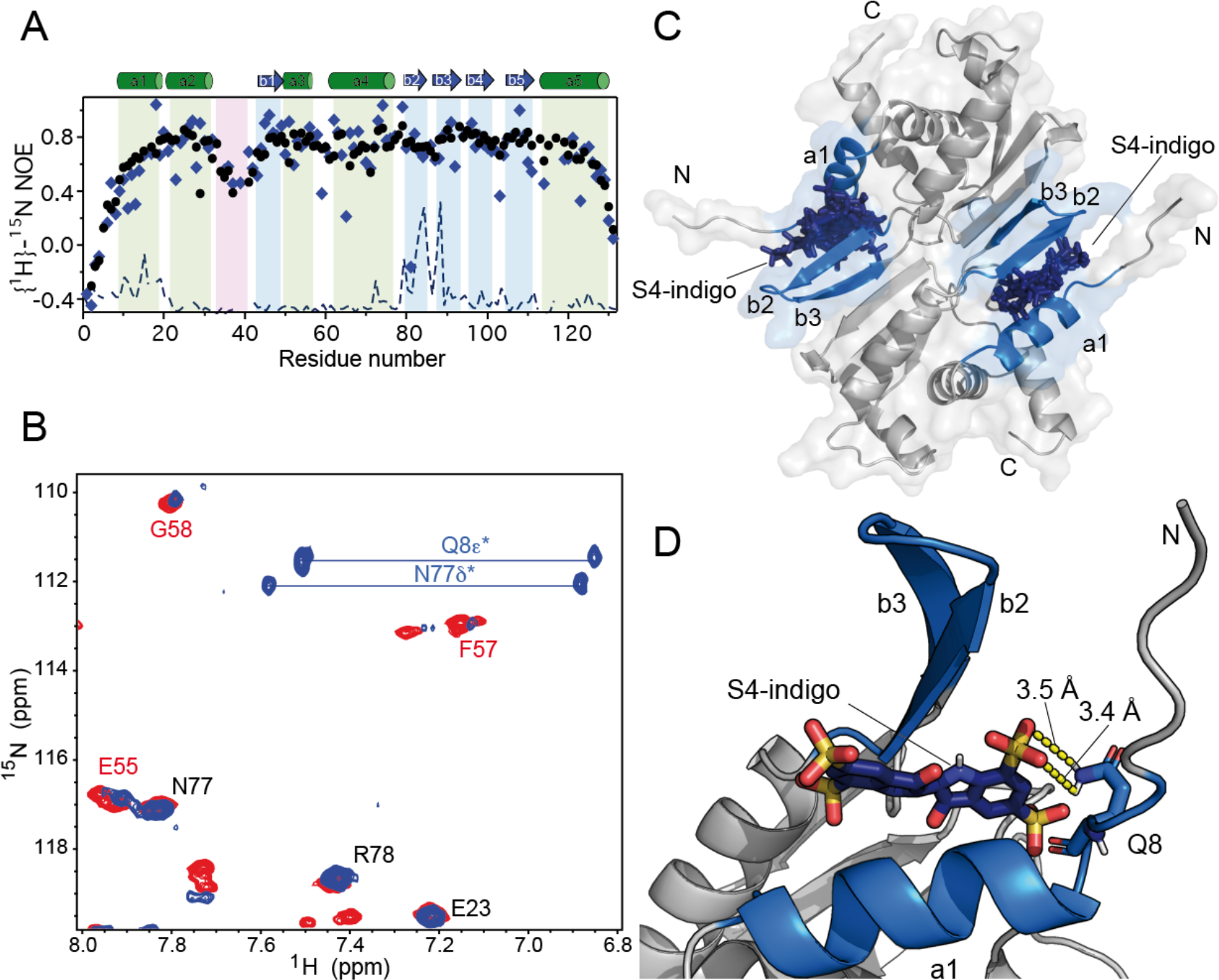
Assessment of S4-indigo Induced Changes in RepA-WH1 Structure and Dynamics. **A.** {^1^H}-^15^N NOE ratios of individual residues in ^13^C,^15^N labeled RepA-WH1. Black dots represent apo RepA-WH1, and blue diamonds represent RepA-WH1 bound to 4 eq. of S4-indigo. The blue dashed line shows the ^1^H-^15^N CSP caused by S4-indigo. **B.** ^1^H-^15^N HQSC spectra of ^13^C,^15^N labeled RepA-WH1 after one night of exchange in 100% D_2_O, 1.0 mM deuterated acetic acid pH* 3.6, 37.0 °C, in the absence (red peaks) or presence (blue peaks) of excess S4-indigo. ^1^H^15^N crosspeaks that are only protected in the absence of S4-indigo have red labels, those only protected in the presence of S4-indigo are labeled blue and those signals which are protected in both conditions have black labels. Astericks refer to peaks with tentative assignments. **C.** Haddock-based rigid-body modeling of S4-indigo bound to RepA-WH1 dimer. α-helix 1 and β-strands 2-3, whose backbone ^1^H^15^N show significant chemical shift perturbations are colored blue, and two ensembles (one per subunit) of ten conformers each of S4-indigo bound to RepA-WH1 are shown in **dark blue**. **D.** Detail of a representative conformer of S4-indigo bound to RepA-WH1 calculated using Haddock based on chemical shift perturbations and H/D exchange derived distance constraints between the sulfonic acid moiety of S4-indigo and the two side chain HNε of Gln 8.

To further monitor alterations in the hydrogen bonding in RepA-WH1 when combined with 4 eq. of S4-indigo, we monitored H/D exchange using NMR spectroscopy (**Sup. Fig. 5**). Although S4-indigo provokes partial aggregation that weakens signal strength, H/D exchange of representative signals from every element of secondary structure could be monitored. The overall pattern of stability is similar relative to RepA-WH1 in the absence of S4-indigo with α-helix 4 and β-strand 4 showing high protection against exchange (**Sup. Fig. 5**). Remarkably, several signals from the β-strands 3, 4 and 5, which form and abut the domain interface; namely V88, W94, F96, V107, L109 and G110, are significantly weaker in the presence of S4-indigo. In contrast, some HN groups preceding or inside α-helix 1 and β-strand 2 (namely Q5, Q8, N10, S16, R82 and V84) show slowed exchange only the presence of S4-indigo. Interestingly, two pairs of slow exchanging side chain ^15^N^1^H_2_ resonances, one from an Asn residue and one from a Gln residue, could be observed in the presence of a four-fold excess of S4-indigo, but not in its absence (Fig. 5B). It is interesting that both side chain amide hydrogens in the asparagine and glutamine residues exchange out at the same rate. Similar results for buried asparagine side chains forming intramolecular contacts in basic pancreatic trypsin inhibitor^61^ have been observed, and are consistent with slow, cooperative unfolding events. There are four Asn and four Gln residues in RepA-WH1. Based on the side chain ^1^H_2_^15^N, ^13^CO, ^13^Cγ, ^13^Cβ and ^13^Cα chemical shift values, the two slow exchanging H_2_N signals could be tentatively assigned to the side chain amide groups of: 1) N4 or N77 and 2) Q8 or Q130. N77’s H_2_Nδ is near S4-indigo at the main binding site ^14^, which strongly suggests that it is protected against H/D exchange in the presence of S4-indigo.

The program HADDOCK and the chemical shift perturbations (CSP) were utilized to calculate a atomistic structural model for S4-indigo united at the secondary binding sites of RepA-WH1. One hundred structures were generated; the fifty lowest energy structures showed an average total energy function of −3500 +/− 60 kcal/mol, which is highly favorable. The thirty structures with the lowest energies were visually evaluated. In all cases, the ligand fit snuggly and filled the gap between α-helix 1 and β-strand 2, and its sulfonic acid moieties are positioned to interact favorably with K11, S15, T18, R81, R82 and Y83 (Fig. 5C). It is also close to Q8 but far from N4, Q5 and N77. Based on these structural models, we tentatively conclude that Q8’s, and not Q130’s, H_2_Nε’s are hydrogen bonded to S4-indigo at these secondary sites (Fig. 5B). The HADDOCK calculations were therefore repeated, using both the CSP data and an indigo-S4-<u>S</u>O_3_^−^ to H_2_<u>N</u>ε distance constraint (3.2 angstrom) as input. Since the resulting structural models place S4-indigo close to Q8 Nε without loss of the other stabilizing contacts and maintain similar energetics (Fig. 5D), these results lend support to the idea that Q8 H_2_Nε could be H-bonded to S4-indigo.

## Discussion

### Differences in the RepA-WH1 solution and crystal structures are pertinent for amyloidogenesis

To assign RepA-WH1, a large dimeric protein, we have used an approach based on combining very low ionic strength, high temperature and the TROSY module to yield good quality 3D spectra suitable for assignment, followed by transfer of the assignments to near physiological conditions by a series of spectra recorded under intermediate conditions. Considering the successful assignment of essentially all backbone and ^13^Cβ resonances, we advance that this approach may be generally useful for studying proteins whose size and oligomerization properties thwart solution NMR spectroscopy in quasi-physiological buffers.

The solution secondary structure of RepA-WH1 is similar to the 3D structure solved by X-ray crystallography, but there are some notable differences that could be relevant for amyloid formation (Fig. 1). In particular, α-helix 1 and α-helix 5 are shorter in solution, as revealed by conformational chemical shifts, as well as ^3^J_HNHA_ coupling constants, relative to the crystal structure. This difference could be attributed to the higher temperature utilized here; namely 50.0 °C, which may tend to promote partial unraveling of weaker elements of secondary structure. Nevertheless, since the thermal denaturation midpoint of the protein is above 90 °C^4−5, 25^, it seems clear that these helices are relatively unstable. Other factors arising from the technical requirements of X-ray crystallography, *e.g.* crystal cooling with liquid nitrogen, crystal packing interactions, the presence of crowding co-solvents, *etc*, could also affect the observed differences.

In the course of this study, several experiments were performed which can detect the presence of secondary structure and intramolecular H-bonds. Of these, the observation of protection against H/D exchange is sensitive to H-bond breaking on slow timescales (Fig. 3). Some HN groups, such as those on the exposed side of the edge β-strands, or in the first turn of α-helices generally do not form H-bonds unless they participate in capping interactions^62^. Thus, a residue could be in an α-helical or β-strand conformation and not show protection against H/D exchange. Despite this consideration, it is safe to conclude that the N-terminal half of α-helix 1 is disordered in solution as revealed by the high ^3^J_HNHA_ coupling constants and the conformational chemical shifts, as these parameters arise physically from peptide bond geometry and not hydrogen bonding. Interestingly, optogenetic devices that modulate the amyloidogenesis of RepA-WH1 upon illumination with blue light, thus generating oligomers cytotoxic in bacteria, have been recently engineered by fusing the N-terminus of the α-helix 1 to the C-terminus of the α-helix J of the *Avena sativa* photoreceptor LOV2^63^. These optogenetic tools open a way to the development of novel light-triggered anti-bacterials (Optobiotics). The results reported here on the enhanced structural dynamics of α-helix 1 provide a rationale for the acute sensitivity of the LOV2-WH1 chimeras to the blue light.

### RepA-WH1’s conformational stability and rigidity are elevated but not uniform

The overall conformational stability of RepA-WH1 is remarkably high, as evinced by the excellent quality and lack of denatured peaks in spectra recorded at 50.0 °C and low pH (Fig. 1A) as well as the elevated H/D protection factors (in excess of 10^6^) at 37.0 °C, pH* 5.7 (Fig. 3). The inter-subunit H-bonds: R91 NH ||| OC V98’, R93 NH ||| OC F98’ and V98 NH ||| OC R91’ strongly protect their HN from exchange. These data suggest that dissociation is strongly coupled to global unfolding, and is consistent with the μM K_D_ values previously reported ^4^. Such strong inter-subunit H-bonding in a β-sheet that spans monomers has never been observed in folded, globular proteins to our knowledge^64^. In other reported H/D exchange studies, the β-strands forming the dimerization interface of globular subunits show weak or little protection such as in RNase H from HIV-1^65^ and the bovine RNase A C-dimer^66^. In stark contrast, the level of H/D protection and conformation stability of intermolecular H-bonds between β-strands can be remarkably high in pathological amyloids such as Aβ ^67^. In the context of the physiological function of RepA-WH1, this is a fascinating finding considering that DNA binding must somehow weaken this tight interface to permit the formation of meta-stable and highly amyloidogenic RepA-WH1 monomers^5^.

In contrast, the first and last α-helices in RepA-WH1 are poor in well-protected HN groups, which implies that these elements of secondary structure are less stable and prone to partial unfolding events on slow (seconds or longer) time scales. It is interesting that α-helix 1, α-helix 5 and the loop following α-helix 2 show considerable dynamic behavior, on ps-ns time scales as gauged by the ^1^H-^15^N relaxation measurements. This intrinsic motion is consistent with the relatively high B-factors seen in these zones by crystallographic analysis^22^ (**Sup Fig. 6**) which was previous viewed as reflecting an inherent tendency or priming to unfold^5^. Our findings concur with that assessment.

### Updated model for RepA-WH1 amyloidogenesis and its inhibition

In addition to the interdomain hydrogen bonds discussed previously^22^, the X-ray structure (PDB: 1HKQ) also reveal an extensive network of cation - π and cation - anion interactions formed by F76, R78, D79, R81, R91, F96 and Y100 which stabilize the dimer interface and communicate with W96 **(Sup. Fig. 7)**. Based on our observations that S4-indigo binding at this site weakens H-bonding at the dimer interface (Fig. 5D) we propose that the selective union of *opsp* DNA, at the dimer interface would sequester Arg and aromatic side chains from these stabilizing interdomain interactions, effectively unlocking the dimerization interface (Fig. 6). These perturbations would then propagate to the helical subdomain via R78 and W94, unleashing the amyloidogenic loop.

**Fig. 6.**
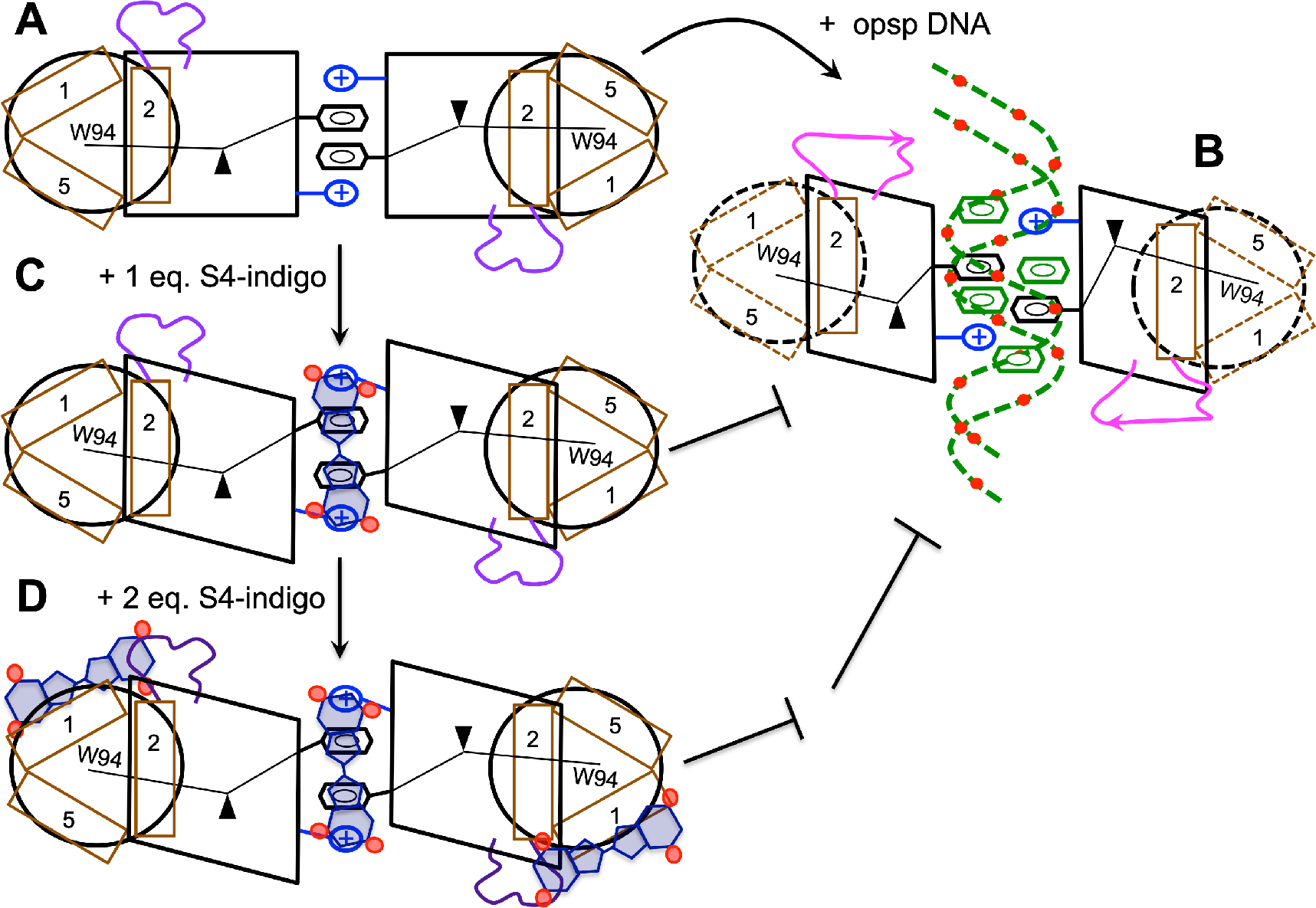
Model for RepA Structural Changes Induced by *opsp* DNA and S4-Indigo Binding. **A.** The intersubunit interface is stabilized by H-bonds as well as a series of cation − π interactions. There is cooperative packing among residues at the interface, in the stable, rigid subdomain formed by β-strands 3, 4 and α-helices 2, 3 and 4 (squares) and in the more dynamic helical subdomain (circles) composed of α-helices 1, 2 and 5 (brown rectangles), with W94 at the heart of its small hydrophobic core. This cooperativity is represented by a fulcrum. The two subdomains are linked by a flexible, amyloidogenic loop (purple). **B.** Binding of *opsp* DNA, with preference over other DNA sequences, at the dimer interface via phosphate (red dots) − arginine charge − charge interactions and specific base (green hexagons) − aromatic residue recognition events unlock the dimerization interface, producing alterations in the hydrophobic core packing and a 30° displacement of the β2-β3 hairpin with respect to the rest of the β-sheet. These perturbations propagate to the helical subdomain, unleashing the amyloidogenic loop (magenta). **C.** The binding of one equivalent of S4-indigo at the subunit interface does not completely unlock the network of cation − π interactions, but does sterically block the binding of *opsp* DNA at the same site. **D.** Addition of two or more equivalents of S4-indigo leads to binding at secondary sites producing contacts among the amyloidogenic loop, α-helices 1 and 5 and the β-hairpin formed by β-strands 2 and 3 which tether the subdomains together, increasing the structure formation in α-helix 1. Note that the β-hairpin formed by β-strands 2 and 3 would lie behind α-helix 1 in the perspective of the cartoon. Thus amyloid formation is blocked not only via steric blocking of binding of *opsp* DNA binding but also by pinning down the amyloidogenic loop (dark purple).

The binding of S4-indigo to secondary sites at a pocket between α-helix 1 and β-strand 2 (two per dimer) in RepA-WH1 suggests an alternative mechanism for the inhibition of amyloid formation (Fig. 6). Whereas the results indicate that the binding of S4-indigo to these secondary sites does not increase the length or stiffness of the terminal α-helices on fast (ps/ns) timescales (Fig. 5A), α-helix 1 and β-strand 2 do show decreased dynamics on slow timescales (Fig. 5B). The tethering of these elements of secondary structure by S4-indigo binding could reduce the local unfolding of the helical subdomain formed by helices 1, 2 and 5 and the loop following helix 2, thereby inhibiting amyloid formation. At higher concentrations of S4-indigo, this mechanism could act in concert with another mechanism previously proposed^14^, in which S4-indigo blocks amyloid formation by sterically impeding the interaction of RepA-WH1 with a specific DNA that triggers amyloid formation. Beside this, by comparing a subunit in the crystal structure of the RepA-WH1 dimer with a structural model of the monomer based on the conformation of the homolog RepE54, the β-hairpin composed by β-strands 2 and 3 was proposed to bend by 30° during RepA-WH1 monomerization^22^. As shown here, binding of S4-indigo to this β-hairpin might prevent this displacement, and thus dissociation and RepA-WH1 amyloidogenesis.

Like RepA-WH1, Transthyretin amyloidogenesis starts from a well folded oligomeric state. Whereas Tafamidis was designed to inhibit Transthyretin amyloidogenesis by stabilizing the native tetramer^68^, our results with RepA-WH1 show that ligands can have multiple effects; namely, 1) Sterically block the union of allosteric effectors like *opsp* DNA, 2) Destabilize the domain interface and 3) Tether flexible loops and partial structured elements to inhibit amyloidogenesis. This rich palette of effects is likely to be general and could be useful to guide the design of future amyloid modulators.

Recently, it has been shown that certain co-chaperones of Hsp90 dichotomously promote healthy refolding or induce the formation of toxic, amyloid-prone conformations of the protein Tau^69^. In *E. coli* cytosol, the Hsp70 chaperone DnaK remodels and detoxifies the amyloid aggregates of RepA-WH1^16^. In a cell-free expression system and in cytomimetic lipid vesicles, the same chaperone also counteracts the amyloid formation of RepA-WH1^26^. In the future, the combination of cell-free expression and the NMR assignments reported here might pave the way for atomistic studies of RepA-WH1/chaperone interactions, which considering the advances obtained from the mechanistic studies of RepA-WH1 cytotoxicity *in vivo*^20^, and could well provide important insights into human neurodegenerative diseases.

## Acknowledgements

This work was supported by grant SAF2016-76678-C2-2-R (DVL), CTQ2017-84371-P (MªÁngeles Jiménez), BIO2015-68730-R (RG) and BFU2015-72271-EXP (RG). NMR experiments were performed in the “Manuel Rico” NMR laboratory (LMR) of the Spanish National Research Council (CSIC), a node of the Spanish Large-Scale National Facility (ICTS R-LRB).

