## Supplementary Information for "Conformational Priming of RepA-WH1 for Functional Amyloid Conversion Detected by NMR Spectroscopy"

**Contents:**

**Sup. Table 1.** Chemical Shifts of RepA-WH1 at 50.0ºC, pH 4.0 in 1.0 mM Acetic Acid. *(related to Figure 1 in the main text)*
**Sup. Table 2.** RepA-WH1 $^{1}$HN Temperature Coefficients at Different Concentrations of S4-Indigo. *(related to Figures 5 & 6 in the main text)*

**Sup. Figure 1.** Levitt-Chothia Topology Diagram of RepA-WH1 (*related to the Introduction and Figure 6 in the main text*)
**Sup. Figure 2.** RepA-WH1 $^{1}$H-$^{15}$N Relaxation and {$^{1}$H}-$^{15}$N NOE *(related to Figure 2 in the main text)*
**Sup. Figure 3**. Participation of Arg Side Chains in S4-Indigo Binding *(related to Figures 4, 5 & 6 in the main text)*
**Sup. Figure 4.** RepA-WH1 $^{3}J_{HNHA}$, δ $^{13}$CO, and {$^{1}$H}-$^{15}$N NOE in the Presence of 4x S4-Indigo. *(related to Figures 4, 5 & 6 in the main text)*
**Sup. Figure 5**. RepA-WH1 H/D Exchange at pH 4.0 Without and With S4-Indigo *(related to Figures 4, 5 & 6 in the main text)*
**Sup. Figure 6**: RepA-WH1 Crystallographic B-factors.
**Sup. Figure 7**. Stabilizing Interactions at the Domain Interface *(related to Figure 6 in the main text)*

**Sup. Table 1.** Chemical Shifts of RepA-WH1 at 50.0ºC, pH 4.0 in 1.0 mM Acetic Acid.

| #NUM | AA | CA | CB | CO | N | HN |
|---|---|---|---|---|---|---|
| 1 | M | 55.91 | 33.01 | 175.61 | 123.30 | 8.71 |
| 2 | V | 62.60 | 32.86 | 175.47 | 121.71 | 8.29 |
| 3 | E | 56.37 | 30.21 | 175.43 | 124.81 | 8.55 |
| 4 | N | 53.28 | 39.10 | 174.62 | 121.13 | 8.60 |
| 5 | Q | 56.87 | 32.91 | 176.25 | 122.37 | 8.46 |
| 6 | V | 63.03 | 32.68 | 175.99 | 121.62 | 8.33 |
| 7 | T | 62.38 | 69.84 | 174.39 | 117.70 | 8.31 |
| 8 | Q | 56.61 | 29.37 | 175.98 | 122.80 | 8.53 |
| 9 | S | 59.37 | 63.69 | 174.23 | 116.92 | 8.50 |
| 10 | N | 54.36 | 38.77 | 175.86 | 121.61 | 8.56 |
| 11 | K | 57.91 | 31.55 | 177.57 | 122.30 | 8.55 |
| 12 | L | 57.40 | 41.80 | 178.12 | 122.10 | 8.27 |
| 13 | I | 63.63 | 37.19 | 178.22 | 120.52 | 8.13 |
| 14 | E | 59.20 | 29.62 | 178.42 | 122.56 | 8.56 |
| 15 | S | 61.27 | 63.91 | 176.31 | 115.47 | 8.47 |
| 16 | S | 61.34 | 63.41 | 174.72 | 118.78 | 8.73 |
| 17 | H | 57.07 | 29.03 | 175.30 | 0.00 | 0.00 |
| 18 | T | 62.28 | 70.19 | 173.75 | 110.81 | 8.13 |
| 19 | L | 55.18 | 43.01 | 177.20 | 121.20 | 7.35 |
| 20 | T | 62.24 | 27.09 | 175.26 | 112.59 | 8.45 |
| 21 | L | 59.32 | 41.56 | 178.42 | 123.46 | 8.91 |
| 22 | N | 56.04 | 37.08 | 177.85 | 114.75 | 9.10 |
| 23 | E | 60.73 | 32.83 | 177.17 | 119.80 | 7.43 |
| 24 | K | 60.86 | 32.75 | 178.00 | 120.17 | 8.62 |
| 25 | R | 59.67 | 30.11 | 177.25 | 117.39 | 8.42 |
| 26 | L | 59.66 | 42.29 | 177.69 | 121.47 | 8.16 |
| 27 | V | 67.66 | 30.90 | 176.98 | 119.57 | 8.45 |
| 28 | L | 58.67 | 41.69 | 179.15 | 121.36 | 8.75 |
| 29 | C | 64.05 | 27.13 | 177.10 | 120.02 | 8.89 |
| 30 | A | 55.38 | 18.57 | 177.74 | 123.35 | 8.81 |
| 31 | A | 54.62 | 18.82 | 179.04 | 118.12 | 8.81 |
| 32 | S | 61.09 | 63.61 | 174.48 | 113.42 | 8.20 |
| 33 | L | 56.07 | 42.68 | 177.43 | 121.78 | 7.80 |
| 34 | I | 61.21 | 37.96 | 174.64 | 119.84 | 7.59 |
| 35 | D | 53.28 | 42.22 | 176.42 | 125.20 | 8.28 |
| 36 | S | 60.22 | 63.82 | 174.32 | 119.22 | 8.57 |
| 37 | R | 56.84 | 30.16 | 175.58 | 120.36 | 8.70 |
| 38 | K | 54.03 | 33.09 | 173.12 | 121.31 | 8.05 |
| 39 | P | 63.13 | 32.49 | 176.49 | 0.00 | 0.00 |
| 40 | L | 53.36 | 41.44 | 174.96 | 124.94 | 8.53 |
| 41 | P | 63.12 | 32.48 | 177.35 | 0.00 | 0.00 |
| 42 | K | 58.27 | 32.28 | 176.81 | 125.06 | 8.76 |
| 43 | D | 55.17 | 40.36 | 174.74 | 120.21 | 8.90 |
| 44 | G | 46.40 | 0.00 | 173.69 | 104.84 | 8.26 |
| 45 | Y | 58.68 | 39.76 | 175.63 | 119.07 | 7.75 |
| 46 | L | 53.87 | 46.74 | 175.27 | 122.43 | 9.19 |
| 47 | T | 62.91 | 69.59 | 172.82 | 120.99 | 8.79 |
| 48 | I | 59.59 | 38.71 | 174.32 | 127.76 | 9.15 |
| 49 | R | 55.64 | 32.03 | 176.80 | 126.93 | 8.89 |
| 50 | A | 55.59 | 18.13 | 178.04 | 125.89 | 9.31 |
| 51 | D | 57.22 | 38.93 | 178.15 | 116.85 | 9.21 |
| 52 | T | 66.30 | 69.33 | 174.85 | 119.64 | 8.00 |
| 53 | F | 61.82 | 40.89 | 176.17 | 123.32 | 8.22 |
| 54 | A | 55.29 | 18.84 | 179.43 | 119.13 | 8.84 |
| 55 | E | 58.78 | 28.62 | 178.50 | 117.39 | 8.15 |
| 56 | V | 65.58 | 32.16 | 177.00 | 118.58 | 8.42 |
| 57 | F | 58.48 | 38.95 | 174.87 | 113.53 | 7.41 |
| 58 | G | 47.26 | 0.00 | 174.35 | 110.76 | 8.03 |
| 59 | I | 60.02 | 41.26 | 174.23 | 118.63 | 7.92 |
| 60 | D | 54.53 | 42.32 | 177.73 | 123.71 | 8.45 |
| 61 | V | 65.98 | 31.95 | 177.28 | 124.41 | 8.80 |
| 62 | K | 58.92 | 31.48 | 177.86 | 120.89 | 8.53 |
| 63 | H | 56.52 | 29.21 | 175.23 | 116.05 | 8.31 |
| 64 | A | 56.15 | 20.10 | 177.74 | 124.71 | 8.18 |
| 65 | Y | 63.41 | 37.40 | 175.98 | 119.10 | 9.10 |

| | | | | | | |
|---|---|---|---|---|---|---|
| 66 | A | 55.15 | 17.68 | 180.08 | 121.90 | 8.32 |
| 67 | A | 54.74 | 18.20 | 178.82 | 120.92 | 7.89 |
| 68 | L | 57.77 | 42.50 | 176.97 | 121.81 | 7.99 |
| 69 | D | 57.37 | 40.76 | 177.78 | 119.70 | 8.75 |
| 70 | D | 58.04 | 42.67 | 176.98 | 120.46 | 8.14 |
| 71 | A | 55.03 | 19.47 | 178.18 | 120.55 | 8.50 |
| 72 | A | 54.99 | 18.77 | 178.15 | 120.02 | 8.91 |
| 73 | T | 67.26 | 69.23 | 175.38 | 115.71 | 8.42 |
| 74 | K | 60.79 | 32.98 | 179.55 | 122.12 | 8.60 |
| 75 | L | 58.01 | 42.55 | 180.04 | 121.66 | 8.76 |
| 76 | F | 61.36 | 39.31 | 176.14 | 120.72 | 8.53 |
| 77 | N | 53.88 | 39.35 | 174.99 | 117.62 | 8.05 |
| 78 | R | 56.19 | 31.95 | 176.99 | 119.27 | 7.65 |
| 79 | D | 52.91 | 44.12 | 173.71 | 123.01 | 9.16 |
| 80 | I | 61.75 | 39.98 | 174.94 | 125.24 | 9.38 |
| 81 | R | 54.64 | 34.59 | 174.55 | 126.60 | 8.84 |
| 82 | R | 55.17 | 34.20 | 174.09 | 121.15 | 8.24 |
| 83 | Y | 57.14 | 42.30 | 176.15 | 126.17 | 9.25 |
| 84 | V | 63.15 | 35.00 | 175.77 | 122.34 | 9.25 |
| 85 | K | 57.56 | 30.36 | 175.95 | 128.91 | 9.76 |
| 86 | G | 45.62 | 0.00 | 173.29 | 104.14 | 8.79 |
| 87 | K | 54.37 | 34.88 | 175.09 | 121.80 | 8.19 |
| 88 | V | 63.51 | 31.87 | 176.03 | 124.61 | 8.77 |
| 89 | V | 61.84 | 33.08 | 175.96 | 120.03 | 8.96 |
| 90 | E | 56.02 | 34.71 | 173.69 | 122.88 | 7.93 |
| 91 | R | 54.70 | 34.31 | 174.21 | 120.98 | 8.92 |
| 92 | M | 54.82 | 38.62 | 173.54 | 123.53 | 9.59 |
| 93 | R | 52.33 | 33.13 | 177.76 | 120.61 | 8.71 |
| 94 | W | 61.27 | 30.62 | 176.21 | 122.73 | 9.92 |
| 95 | V | 57.75 | 35.91 | 173.36 | 106.19 | 7.05 |
| 96 | F | 55.50 | 41.32 | 173.40 | 116.73 | 8.73 |
| 97 | H | 53.83 | 36.93 | 172.73 | 124.35 | 7.19 |
| 98 | V | 60.88 | 35.55 | 172.70 | 126.95 | 9.48 |
| 99 | K | 54.91 | 34.48 | 174.03 | 125.83 | 8.95 |
| 100 | Y | 57.99 | 38.42 | 175.48 | 125.28 | 8.83 |
| 101 | R | 53.66 | 29.16 | 175.66 | 127.66 | 8.56 |
| 102 | E | 60.26 | 29.18 | 178.94 | 127.04 | 8.70 |
| 103 | G | 46.49 | 0.00 | 174.60 | 108.09 | 9.11 |
| 104 | Q | 55.35 | 30.19 | 175.69 | 117.62 | 7.53 |
| 105 | G | 47.40 | 0.00 | 173.96 | 109.83 | 8.16 |
| 106 | C | 56.29 | 32.40 | 171.33 | 114.55 | 8.40 |
| 107 | V | 60.33 | 35.36 | 172.94 | 114.99 | 8.62 |
| 108 | E | 54.20 | 31.71 | 174.37 | 124.96 | 9.11 |
| 109 | L | 53.46 | 45.69 | 174.16 | 124.86 | 9.30 |
| 110 | G | 45.32 | 0.00 | 172.61 | 110.53 | 8.87 |
| 111 | F | 59.09 | 40.43 | 174.81 | 126.43 | 9.06 |
| 112 | S | 57.40 | 63.42 | 173.54 | 118.27 | 8.57 |
| 113 | P | 65.51 | 31.93 | 176.84 | 0.00 | 0.00 |
| 114 | T | 62.39 | 68.99 | 174.89 | 103.96 | 7.12 |
| 115 | I | 60.34 | 38.52 | 176.62 | 121.25 | 8.01 |
| 116 | I | 65.78 | 35.96 | 173.85 | 120.67 | 7.43 |
| 117 | P | 65.41 | 31.15 | 176.92 | 0.00 | 0.00 |
| 118 | H | 55.85 | 30.32 | 175.77 | 114.11 | 7.93 |
| 119 | L | 58.70 | 41.88 | 178.15 | 0.00 | 0.00 |
| 120 | T | 66.03 | 68.67 | 176.33 | 113.07 | 8.55 |
| 121 | M | 58.14 | 32.40 | 178.05 | 122.67 | 7.84 |
| 122 | L | 57.38 | 41.35 | 177.40 | 120.66 | 8.15 |
| 123 | H | 58.81 | 28.58 | 175.72 | 117.84 | 8.45 |
| 124 | K | 58.57 | 32.56 | 177.58 | 118.87 | 7.90 |
| 125 | E | 58.05 | 29.26 | 177.15 | 119.49 | 8.12 |
| 126 | F | 57.38 | 39.09 | 175.87 | 117.75 | 8.29 |
| 127 | T | 63.21 | 70.24 | 174.75 | 112.49 | 8.00 |
| 128 | S | 59.88 | 63.71 | 173.90 | 117.27 | 8.21 |
| 129 | Y | 57.99 | 38.47 | 174.99 | 120.96 | 7.95 |
| 130 | Q | 56.10 | 29.28 | 175.03 | 120.67 | 8.25 |
| 131 | L | 54.99 | 42.46 | 175.15 | 124.10 | 8.30 |
| 132 | K | 57.57 | 33.80 | 179.79 | 126.92 | 7.90 |

### **Supplementary Table 2:**

Temperature Coefficients at Different Concentrations of S4-Indigo

| Sequence # | 0 Eq. S4-indigo | 4 Eq. S4-indigo | 7 Eq. S4-indigo |
|---|---|---|---|
| 1 | -6.9 | | |
| 2 | -6.9 | | |
| 3 | -6.5 | -7.1 | |
| 4 | -7.2 | -7.6 | |
| 5 | -6.7 | | |
| 6 | | | |
| 7 | -6.6 | -6.9 | |
| 8 | -6.8 | -6.4 | |
| 9 | -5.8 | -5.9 | |
| 10 | -4.7 | -4.6 | |
| **11** | **-4.6** | **-2.5** | |
| 12 | -2.0 | | -3.6 |
| 13 | -5.3 | -4.7 | |
| 14 | -3.5 | -8.6 | |
| 15 | -4.3 | -5.0 | |
| **16** | **-8.2** | **-2.0** | |
| 17 | | | |
| 18 | -2.0 | -1.8 | |
| **19** | **-1.7** | **-2.4** | **0.60** |
| 20 | -1.4 | | |
| **21** | **-4.9** | **-0.50** | |
| **22** | **-6.8** | | **-1.8** |
| 23 | -0.30 | -2.7 | |
| **24** | **-5** | **0.0** | |
| 25 | -1.2 | -6.8 | |
| 26 | -2.7 | -2.9 | -2.6 |
| 27 | -1.7 | -4.1 | |
| 28 | -7.5 | | |
| 29 | -1.0 | -3.1 | |
| **30** | **-2.0** | **-4.9** | |
| 31 | -4.2 | | |
| **32** | **-2.3** | **-6.4** | **-7.2** |
| 33 | -0.50 | | -1.4 |
| 34 | -0.70 | | |
| **35** | **0.72** | **-8.7** | **-7.3** |
| 36 | -10.3 | | |
| 37 | -6.4 | | |
| 38 | -3.0 | | |
| 39 | | | |
| **40** | **-8.1** | **-0.90** | |
| 41 | | | |
| 42 | -8.1 | -8.3 | |
| 43 | -2.1 | | |
| 44 | -3.4 | -3.4 | -4.3 |
| 45 | 1.2 | -2.5 | |
| 46 | -3.0 | -0.90 | |
| 47 | -1.5 | | |
| 48 | -3.5 | -2.8 | |
| **49** | **-6** | **-3.4** | |
| 50 | -1.1 | -2.8 | |
| 51 | -4.8 | -6.2 | -5.5 |
| 52 | -7.5 | | |
| 53 | -3.2 | | |
| 54 | -1.6 | -2.8 | |
| 55 | -1.5 | -2.1 | |
| 56 | -6.3 | -7.1 | |
| 57 | -1.2 | -1.5 | -0.50 |
| 58 | -0.10 | -0.90 | |
| 59 | -1.4 | 0.30 | |
| 60 | -6.6 | -6.9 | |
| 61 | -6.3 | -7.2 | |
| 62 | -4.3 | -3.8 | |
| 63 | -3.8 | | |
| 64 | -2.0 | 1.8 | 1.6 |
| 65 | -6.2 | -7.2 | -6.9 |
| 66 | -9.0 | | |
| 67 | -0.7 | -0.2 | |
| 68 | -1.9 | | |
| 69 | -3.4 | | |
| 70 | -1.6 | | |
| 71 | -1 | | |
| 72 | -4.7 | | |
| 73 | -6.1 | -6.2 | |
| **74** | **-1.7** | **-5.6** | |
| 75 | -2.1 | -2.8 | |
| 76 | -0.50 | -2.6 | |
| 77 | -0.80 | -1.3 | |
| 78 | -0.70 | -1.6 | |
| 79 | -7.3 | -10.2 | -8.9 |
| 80 | -5.6 | -3.7 | -8 |
| 81 | -6.2 | -6.6 | |
| **82** | **-1.1** | **-6.6** | |
| 83 | -7.7 | -7.7 | |
| 84 | -2.8 | | -4.2 |
| 85 | -9.3 | -9.5 | -10.2 |
| 86 | -7.9 | -8 | -7.9 |
| 87 | -1.4 | | |
| 88 | -7.3 | | |
| 89 | -4.8 | -6.3 | |
| 90 | -1.7 | -1.8 | |
| **91** | **-1.9** | **-5.3** | |
| 92 | -0.10 | 1.2 | 0.80 |
| 93 | -3.0 | -2.6 | |
| 94 | -8.4 | -10.3 | -9.6 |
| 95 | -1.9 | 1.7 | |
| 96 | -1.6 | | |
| 97 | -0.1 | -1.2 | -1.3 |
| 98 | -3.0 | -1.5 | -3.2 |
| 99 | -2.9 | | |
| 100 | -1.2 | -1.3 | |
| 101 | -2.6 | -3.1 | -3.5 |
| 102 | -5.9 | -6.8 | -6.8 |
| 103 | -4.5 | -4.8 | -4.4 |
| 104 | -1.4 | -2.5 | -1.3 |
| 105 | -1.9 | -2.5 | -2.7 |
| 106 | -3.5 | -4.1 | -3.8 |
| 107 | -2.2 | -5.7 | |
| 108 | -3.8 | -3.3 | -4.0 |
| 109 | -3.8 | 1.5 | -5.2 |
| 110 | -3.0 | -5.7 | -3.0 |
| 111 | -3.7 | -0.20 | |
| **112** | **-1.5** | **-5.0** | |
| 113 | | | |
| 114 | -2.3 | -0.60 | |
| **115** | **-0.30** | **-7.6** | |
| 116 | | -0.50 | |
| 117 | | | |
| 118 | -2.5 | -6.3 | -6.0 |
| 119 | | | |
| 120 | | -7.6 | -4.4 |
| 121 | -5.1 | -4.8 | -4.8 |
| 122 | -3.4 | -5.5 | |
| 123 | -6.4 | -6.4 | |
| 124 | -1.5 | -2.1 | |
| 125 | -2.1 | -0.7 | |
| 126 | -6.1 | | |
| 127 | -0.9 | -2.5 | -1.6 |
| 128 | -3.6 | -3.7 | -3.1 |
| **129** | **-0.9** | **-4.5** | |
| 130 | -5.9 | | |
| 131 | -7.1 | | -7.3 |
| 132 | -4.1 | -5.9 | -5.8 |

In **Supplementary Table 2**, residues adopting α-helices and β-strands are shaded blue and green, respectively. The amyloidogenic segment is shaded magenta. Residues whose $^{1}$HN temperature coefficients significantly increase (indicating the formation of H-bonds) are written in bold **blue**; on the contrary, those whose $^{1}$HN temperature coefficients drop (which evidences loss of H-bonding) are shown in bold **red**.

**Sup. Figure 1:** Levitt-Chothia Topology Diagram of RepA-WH1

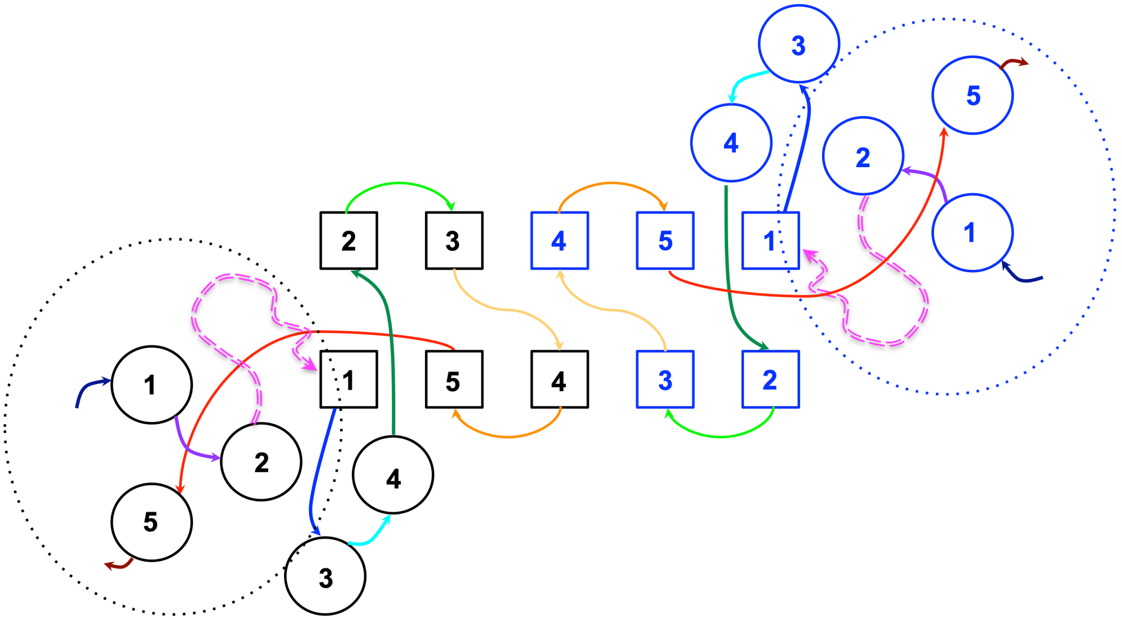


In this 2D diagram of the RepA-WH1 structure, α-helices and β-strands are represented by circles and squares, respectively, which are numbered in order starting at the N-terminus. The connections between the elements of secondary structure are shown as arrows, colored in the order of rainbow from the N-terminus (indigo) to the C-terminus (red), except for the amyloidogenic loop (magenta broken arrow) which connects α-helix 2 and β-strand 1. The helical subdomain formed by α-helices 1, 2 and 5, whose peripheral topology is highly unusual, is highlighted with a dotted circle. The three helix bundle composed of α2, α3 and α4, together with the "wing" made by β1, β5 and β4, constitute the canonical "winged helix" domain fold. α-helix 2 is special; it precedes the amyloidogenic loop and is common to both the periperal (α1, α2, α5) and canonical (α2, α3, α4) three helix subdomains. β-strands 2 and 3 are also distinct from the canonical "winged helix" fold and play key roles in RepA-WH1's dimerization [See Giraldo *et al.,* (2003) Nat. Struct. Biol. **10**(7): 565-571 for more details].

**Sup. Figure 2:** RepA-WH1 $^{1}H$-$^{15}N$ Relaxation and $\{^{1}H\}$-$^{15}N$ NOE

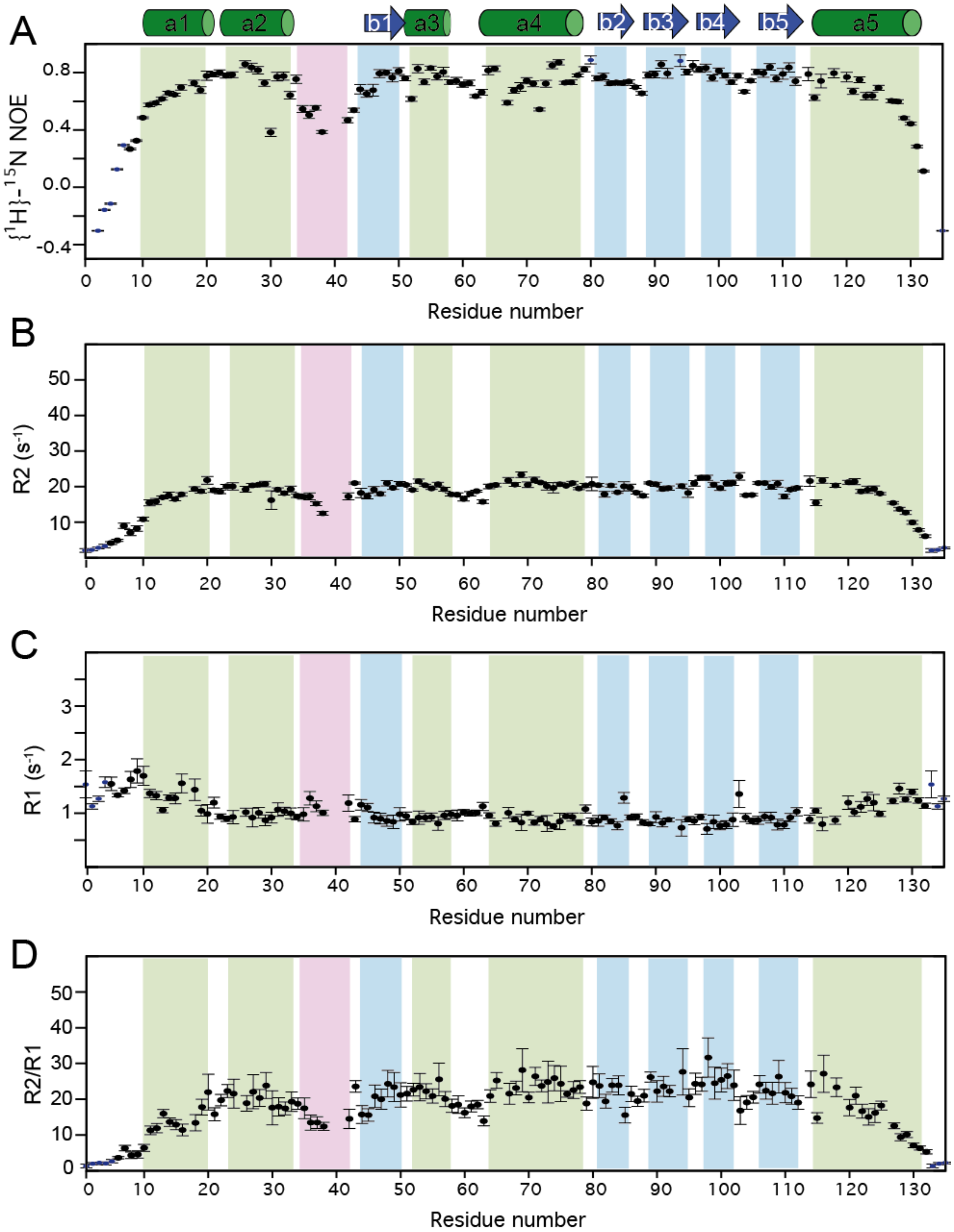


These data provide the basis for the "Model Free" analysis of the dynamics of RepA-WH1 which are shown in **Fig. 2** in the main text. Sample conditions: 1.0 mM DAc ($d_4$), pH 4.0, 50.0 ºC, 85% $H_2O$ mQ, 15 % $D_2O$.

**Sup. Figure 3**: Participation of Arg Side Chains in S4-Indigo Binding.

**A**. Spectra of the upfield "Arg $^{15}N\varepsilon$" region of two $^{1}H$-$^{15}N$ HSQC spectra are shown. Peaks with different degrees of folding were revealed by registering these spectra with distinct $^{15}N$ sweep widths. When suitably aligned along the y-axis, the backbone HN peaks appear as superimposed **blue** and **black** peaks, while side chain Trp, Lys and Arg HN moieties (labeled) can be readily identified as they appear as separated **blue** and **black** peaks.

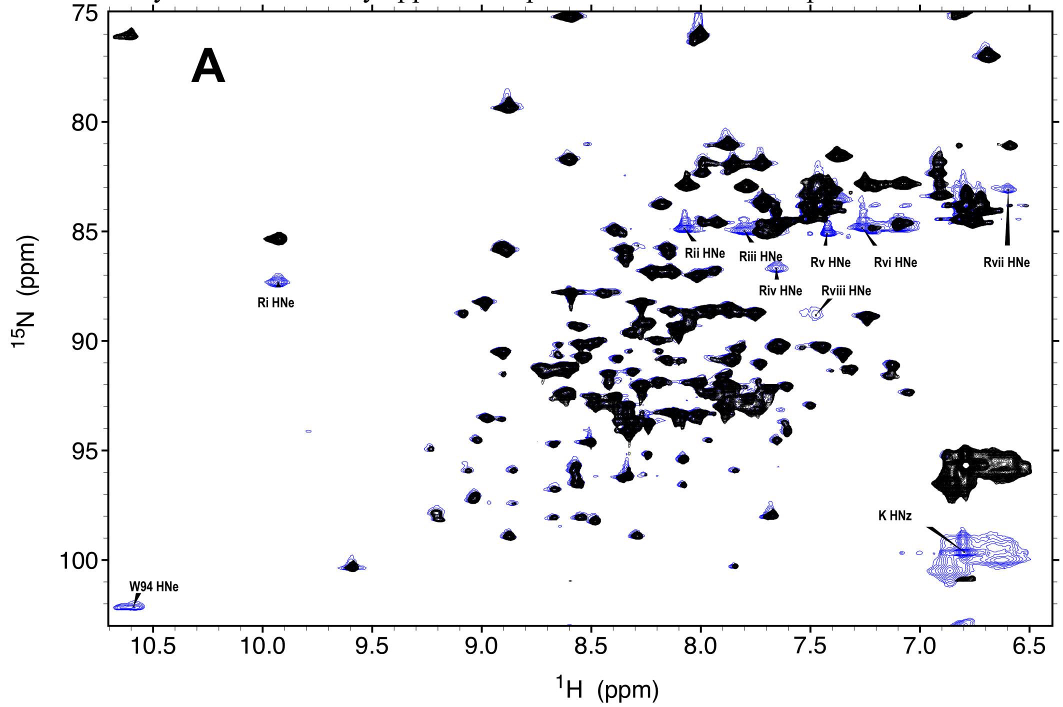


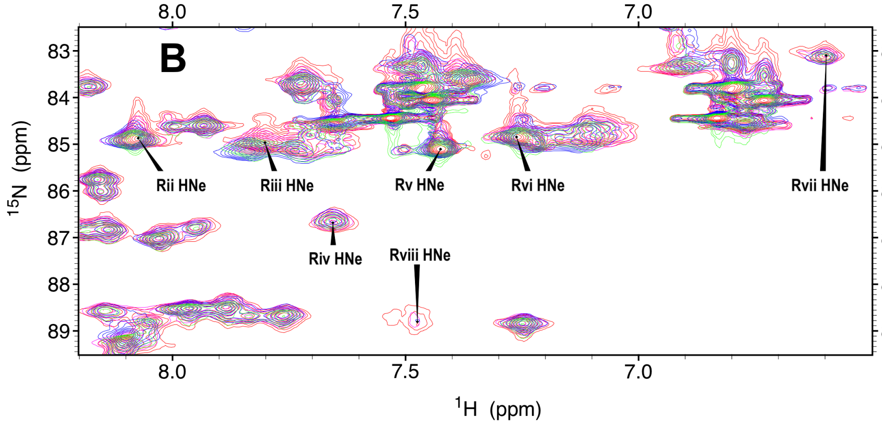


**B & C**. Changes in the Arg HNε peaks in the presence of 0.0 (**red**), 0.4 (**magenta**), 1.0 (**green**) and 2.0 (**blue**) equivalents of S4-indigo per dimer of RepA. Significant chemical shifts changes are observed for the Ri and Riii Arg HNε resonances. The Rviii HNε peak seems to broaden and disappear upon addition of S4-indigo. One Arg side chain resonance (labeled "Ri" see panel **C**) has a significantly downfield shifted $^{1}$HNε resonance (panel **C**) which strongly suggests that it is H-bonded. A survey of the 3D structure (**PDB-1HKQ**) putatively assigns this signal to Arg 78, as this residue's guanidinium group is H-bonded to Glu 23's carboxylate group.

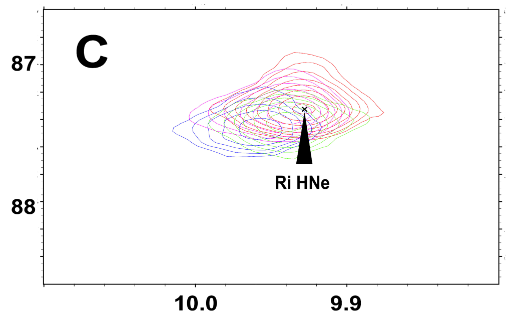

**Sup. Fig. 4:** Changes in the RepA-WH1 Coupling Constants, $^{13}$CO Chemical Shifts and $\{^1H\}$-$^{15}$N NOE Ratio in the Presence of a Four-Fold Excess of S4-Indigo.

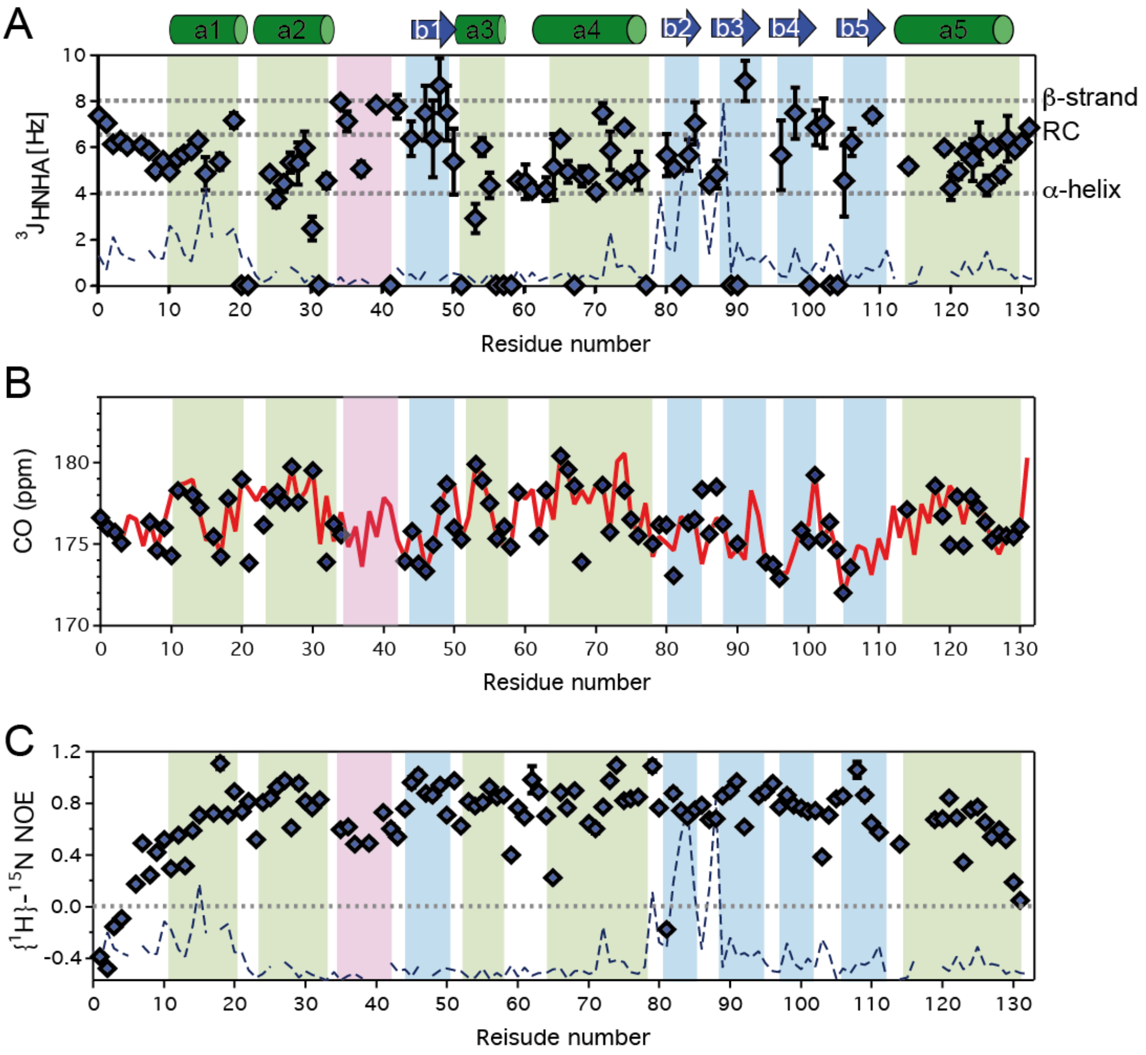


**A.** The $^3J_{HNHA}$ coupling constant, which is indicative of secondary structure, is shown for individual residues in RepA-WH1. No significant trends towards increased secondary structure content is detected in the presence of a four fold molar excess of S4-indigo. The dashed line (labeled CSP) shows the $^1H$-$^{15}N$ chemical shift perturbation in the presence of S4-indigo. Values at zero were close to the noise level. Error bars were calculated based on spectral signal/noise.

**B**. The $^{13}$CO chemical shifts for RepA-WH1 alone (**red** line) or in the presence of 4x excess S4-indigo (**black** diamonds) are quite similar, which is consistent with no changes in secondary structure.

**C**. $\{^1H\}$-$^{15}$N NOE ratios of individual residues in RepA-WH1. The dashed line (CSP) represents the $^1H$-$^{15}N$ chemical shift perturbation in the presence of S4-indigo. Error bars were calculated based on spectral signal/noise.

**Sup. Fig. 5:** RepA-WH1 H/D exchange at 37.0 ºC, pH* 3.6 in the Absence (**A**) or Presence (**B**) of 4x S4-Indigo.

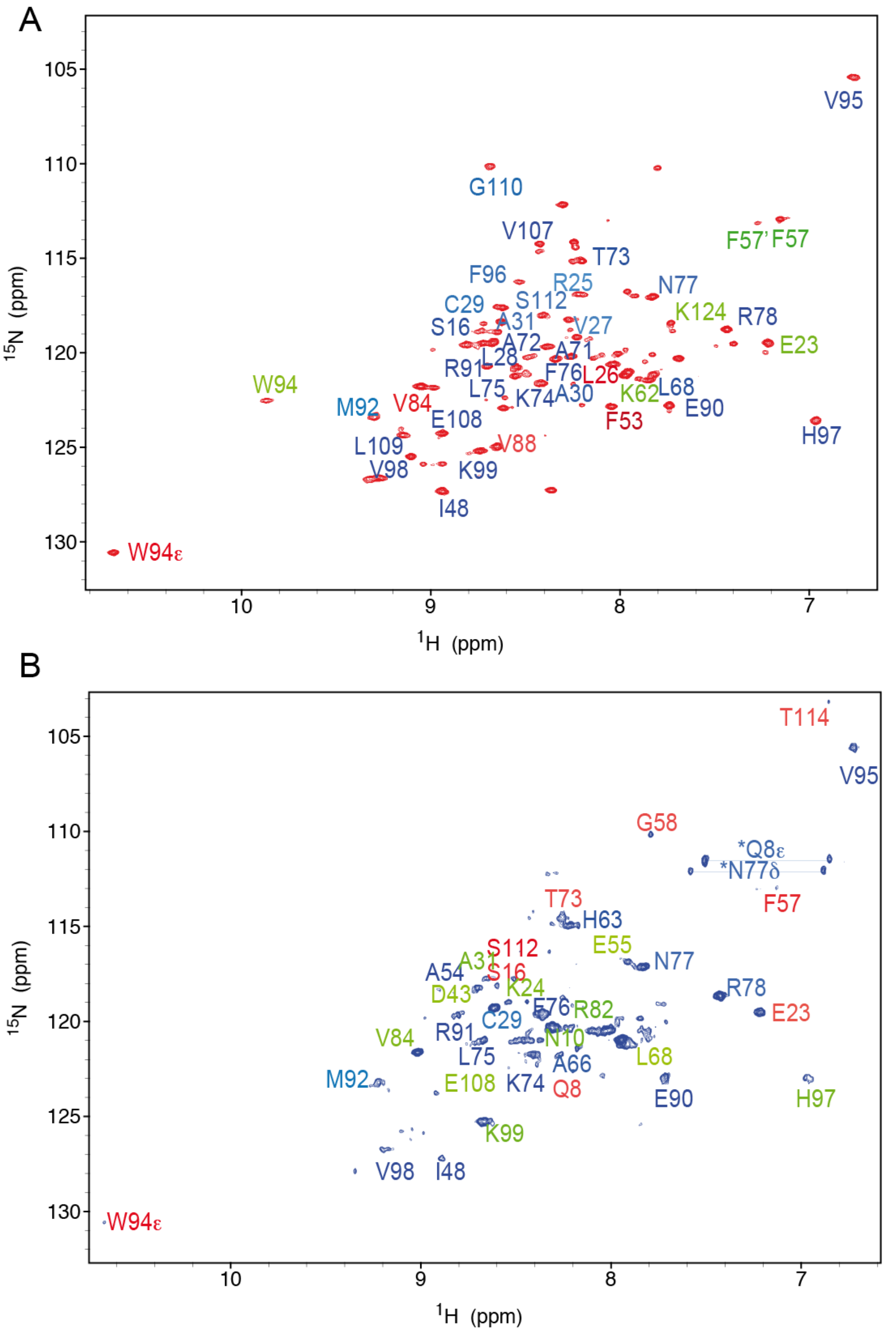


$^{1}H$-$^{15}N$ HSQC spectra of RepA-WH1 domain undergoing H/D exchange at 50.0 ºC in 1.0 mM DAc (d4) and 100 % $D_2O$ pH* 3.6 in absence (**A**) or presence (**B**) of 4x S4-indigo. After recording initial 2D HSQC spectra, 3D HNCO spectra were recorded. The peaks whose identity could not be confirmed by the matching the $^{13}CO$ chemical shift in 3D HNCO experiment are left unlabeled. HNs which are initially protected (after 1.5 hours of exchange) but that exchange after overnight incubation are labeled in **red**. Those resisting exchange for >15 h, but not 30 days are labeled in **green**. Finally HN resisting exchange >30 days are labeled in **blue**. The slow exchanging Asn and Gln side chain $H_2N$ groups are putatively assigned to Asn77 and Gln 8 and the uncertain nature of these assignments is represented by the asterisks.

**Sup. Fig. 6**: RepA-WH1 Crystalographic B-factors.

Two views of one RepA-WH1 monomer from the crystal structure (PDB 1HKQ) with the chain colored in a rainbow series for low B-factors (cyan) to high B-factors (reddish orange). It is notable that the other subunit (not presented here) shows much less variation in its B-factors, which might be due to crystal packing interactions.

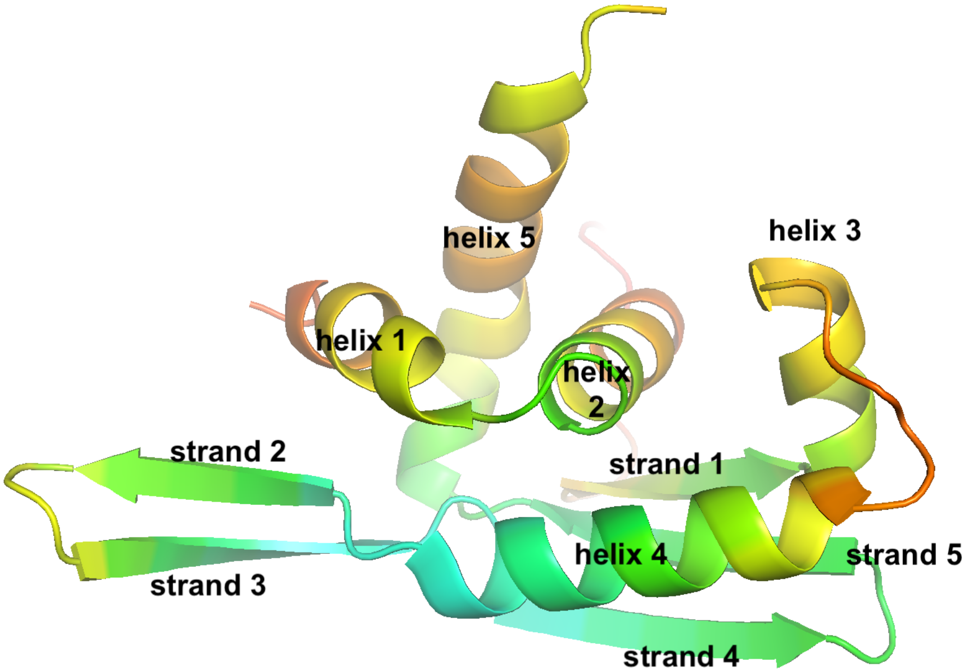


*(rotate 90º about the z-axis)*

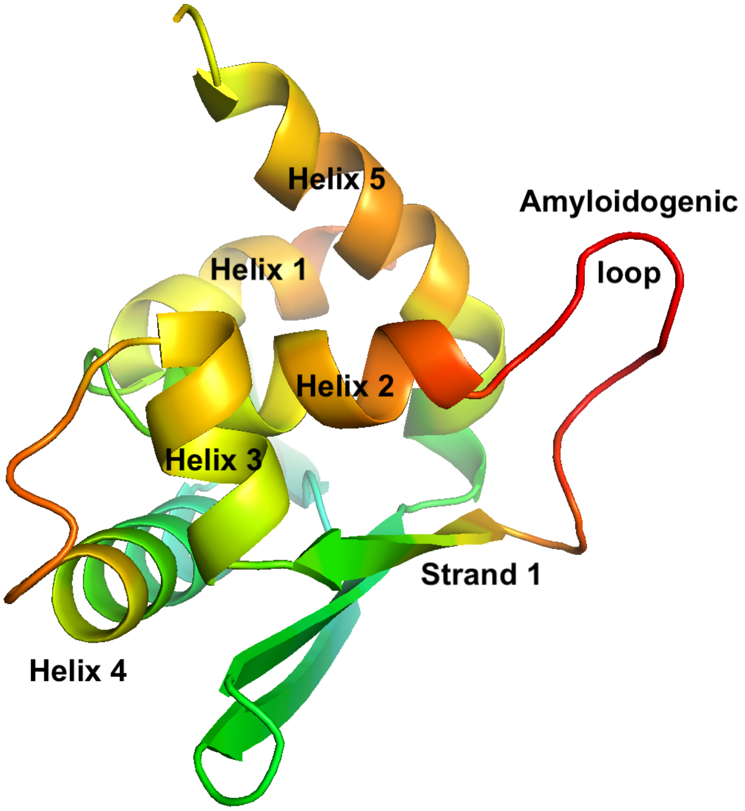

**Sup. Figure 7:**

Stabilizing interactions at the Domain Interface

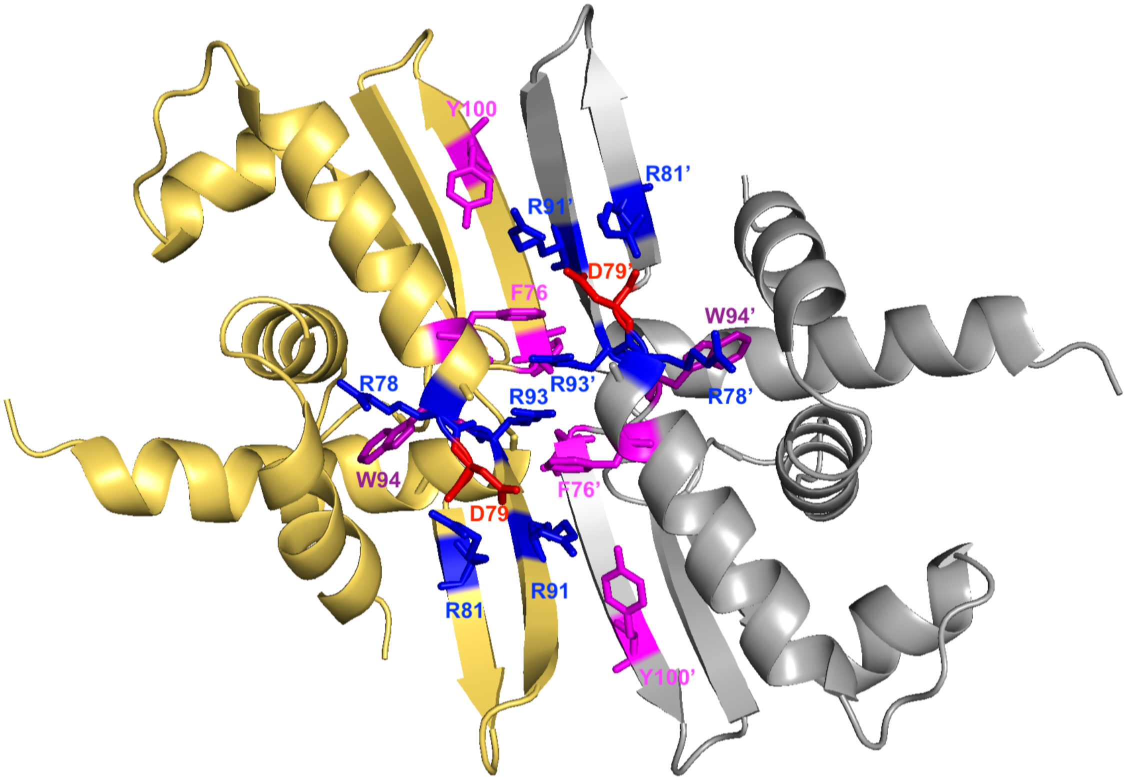


Ribbon diagram of RepA-WH1 showing the two domains in **gold** and **silver**. Aromatic (**magenta**) and charged Arg (**blue**) and Asp (**red**) residues that form stabilizing interdomain interactions are labeled. Trp 94 (**purple**), which is connected to those stabilizing interactions through Arg 78, is also labeled. On the opposite site of the dimer (behind R93), there is a $\pi$ - $\pi$ contact formed by the two Phe 96 sidechains (one from each sidechain, not labeled).
